# Resonance-driven enhancement of sleep spindles using thalamic temporal interference stimulation

**DOI:** 10.64898/2026.05.29.728879

**Authors:** M. Gabriela Navas-Zuloaga, Tobias Raufeisen, Prince Okyere, Esra Neufeld, Niels Kuster, Derk-Jan Dijk, Ines R. Violante, Maxim Bazhenov, Ullrich Bartsch

**Author notes:** Authors contributed equally (M. Gabriela Navas-Zuloaga, Tobias Raufeisen). Co-senior (Maxim Bazhenov, Ullrich Bartsch).

## Abstract

Sleep spindles are hallmarks of non-rapid eye movement sleep and support memory consolidation yet remain difficult to modulate non-invasively.

Combining computational modeling and human sleep recordings, we show that thalamus-targeted temporal interference stimulation (TIS) with a 5Hz envelope increases spindle density via subthreshold resonance in thalamocortical relay neurons.

Our results demonstrate a mechanistic framework for the rational design of interventions to selectively augment sleep spindles.

## Main Text

Sleep spindles are transient waxing-and-waning 9-16Hz brain oscillations originating in the thalamus^1^, characteristic of non-rapid-eye-movement (NREM) sleep and playing a key role in memory consolidation and learning^2^. Spindle deficits are prominent in aging^3^, Alzheimer’s disease^4^ and schizophrenia^5^.

Existing clinical interventions that augment spindle activity rely on pharmacology^6^, but carry significant side effects^7^. Non-invasive brain stimulation (NIBS) using either sound or electrical stimulation have been investigated^8,9^, but were mainly limited to cortical targets, and hampered by inconsistency across studies, with stimulation parameters often chosen heuristically^10^. Targeting spindle circuits more precisely may require stimulation parameters grounded in the biophysical properties of the neurons that generate them.

Spindles arise from reciprocal interactions between thalamocortical (TC) relay neurons and inhibitory thalamic reticular (RE) neurons^1,11,12^. At each spindle cycle, RE bursts hyperpolarize TC neurons, triggering rebound TC bursting mediated by low-threshold (T-type) calcium channels, which in turn recruits further RE activity. Spindle termination is mediated by Ca^2+^-dependent h-current upregulation, which depolarizes TC neurons and limits rebound bursting^13,14^, and network desynchronization^15^. Given that excitability of thalamic neurons is central to spindle initiation and timing we hypothesized that stimulation tuned to intrinsic timescales could be leveraged to selectively engage spindle-generating circuits.

Direct thalamic targeting is now possible non-invasively using temporal interference stimulation (TIS) which applies two high-frequency alternating currents whose interference generates a low-frequency envelope that is demodulated by neurons^16^. The spatial extent of the interference region can be designed to selectively target subcortical regions^16–18^.

Here, we combine biophysically detailed computational modeling with human TIS-EEG recordings during sleep to show that intrinsic subthreshold resonance of TC neurons determines the stimulation frequencies most effective for spindle generation, providing a mechanistic framework for targeted non-invasive modulation of sleep spindles.

To investigate whether subthreshold stimulation can elicit spindle events, we employed a thalamocortical network model of cortical pyramidal (PY), inhibitory (IN), thalamic RE and TC neurons with Hodgkin–Huxley-type dynamics (Figure 1a). The model, extending prior work^19^, generated spontaneous sleep spindles originating in the thalamus and propagating to cortex (Figure 1b). We modeled TIS as a subthreshold sinusoidal current applied to thalamic neurons (Figure 1c). To isolate thalamic mechanisms, corticothalamic projections were initially removed.

**Figure 1.**
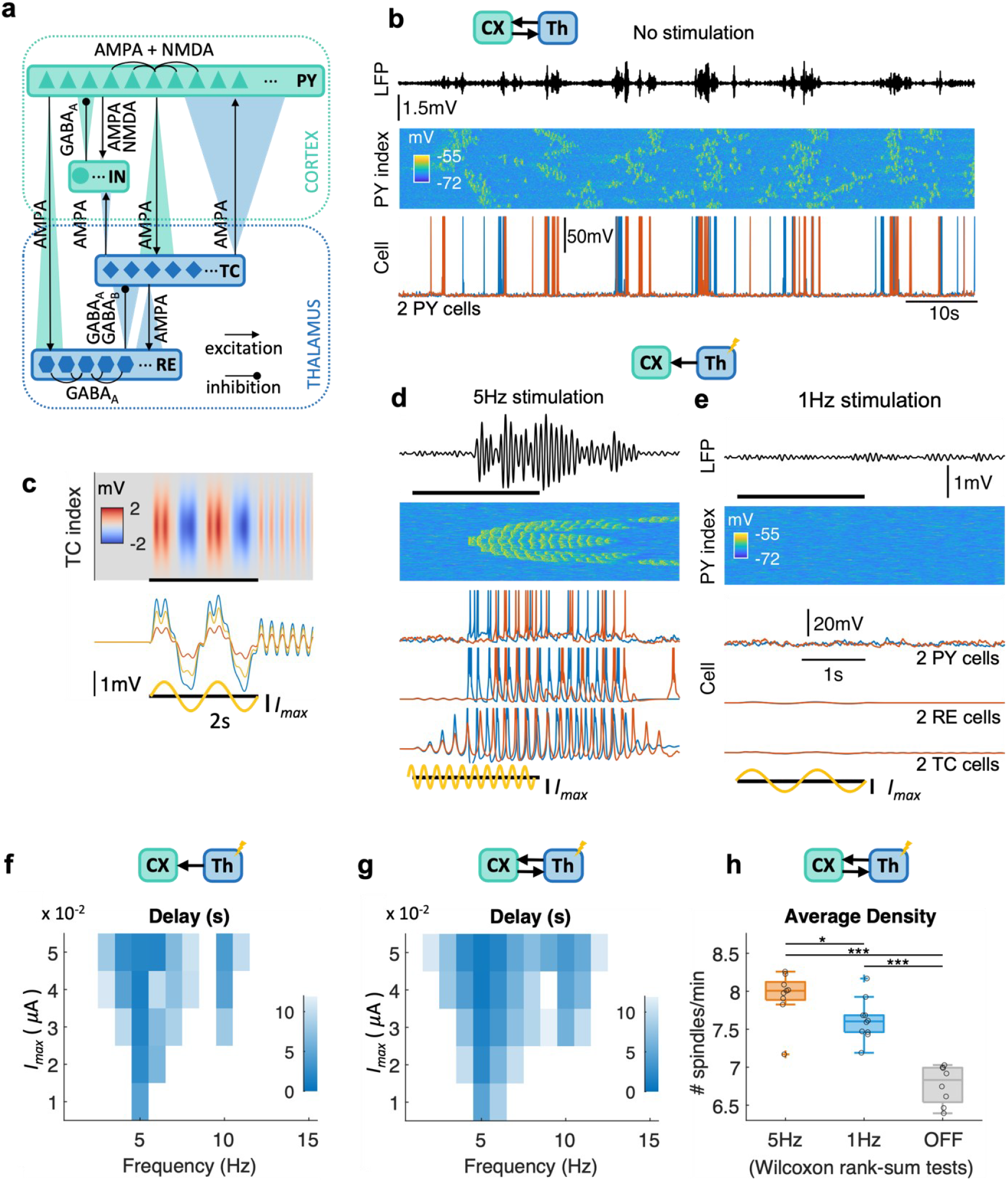
Computational model results (a) Computational model diagram with connections between 1000 pyramidal (PY), 200 inhibitory (IN), 500 thalamocortical (TC), and 500 reticular (RE) populations. (b) Baseline spindle dynamics in cortical LFP (top), PY cell voltages (middle) and two PY cell traces (bottom). (c) Cortical (CX) inputs to the thalamus (Th) are removed to isolate the effect of thalamic stimulation (top). TIS is modeled as sinusoidal current input to TC and RE cells. Heatmap shows voltage deviation from resting potential for TC cells subject to 2s 1Hz sub-threshold stimulation with maximal intensity I_max_ = 0.05 μA (blackbar, sine wave). Linesshow voltage traces for 3 TC cells. Stimulus intensity is maximal at the network center and decays with distance as a gaussian. Injected current generates less than 2mV voltage deviations from resting potential. (d) No spindles elicited for simulated TIS at 1Hz envelope for 2s (black bar, sine wave). From top to bottom: cortical LFP, PY cell voltage heatmap, voltage traces for 2 PY, 2 RE, and 2 TC cells. (e) Spindle in response to 5Hz stimulation. Panels are as (d). Note that TC subthreshold oscillations initiate the spindle cycle. (f) Sweep over stimulation intensity (I_max_ in the range of 0.01-0.05 μA) and frequency (1-15Hz) for the model without cortical inputs to the thalamus. Simulations are 40s with a 2s stimulation period. Color indicates delay between stimulation start and spindle onset. No spindles are detected outside the colored region, which exhibits and Arnold-tongue shape. Delay is minimal across intensities for ~5Hz frequency. (g) As (f), for the model with cortical inputs to the thalamus. The Arnold-tongue region persists, with spindles detected outside only after large delays corresponding to spontaneous inter-spindle intervals. (h) Spindle density for full-model simulations (610s each) without stimulation (OFF) compared to 5Hz or 1Hz stimulation intervals of 2s every 8s (6s refractory period), with I_max_ = 0.03 μA. Asterisks mark significant comparison between density medians from Wilcoxon rank-sum tests between conditions (*p<0.05, ***p<0.001).

For a fixed stimulation amplitude, the modelled spindle responses depended on frequency. Stimulation at 5Hz induced a gradual buildup of subthreshold oscillation, which turned suprathreshold and triggered spindle events, whereas 1Hz stimulation did not (Figure 1d,e). A systematic sweep across frequency–amplitude space revealed an Arnold tongue structure (Figure 1f), with stimulation most effective near 5Hz (lowest latency to spindle onset). Within this region, spindle amplitude and frequency remained stable across stimulation parameters (Figure S1). Reintroducing corticothalamic synaptic projections reproduced these results (Figure 1g).

Repeated 5Hz stimulation every 8s in the full network model increased spindle density compared to 1Hz or no stimulation (OFF) (Figure 1h). 5Hz stimulation robustly generated spindles aligned to stimulation onset when targeting TC neurons alone or TC together with RE neurons (Figure S2). In contrast, 1Hz or 5Hz stimulation of RE, PY, or IN neurons alone, or PY and IN together, produced little or no alignment between stimulation and spindle events (Figure S2), indicating that frequency selectivity arises from specific biophysical properties of TC neurons rather than nonspecific network activation.

To test the model predictions, we performed a nap study in 20 adult participants (mean age=27). TIS electrode locations (Figure 2a) were chosen based on FEM modelling with bilateral EM field exposure optimization for the whole thalamus (Figure S4b). Stimulation was delivered in 8-second blocks with a 1s PRE window, 2s ON, and 5s POST window at df=5Hz and df=1Hz (cf=2000Hz, 2mA peak-to-baseline), and df=0Hz, 0mA OFF condition (Figure 2b). A comparable number of stimulation trials at 5Hz and 1 Hz conditions was delivered during sleep (Tables S1, S2), with a small subset of OFF trials excluded due to switching artifacts during ramping (see Methods). The number of stimulation blocks delivered per sleep stage did not differ between participants. We restricted our analysis to N2 sleep to match the model preconditions.

**Figure 2.**
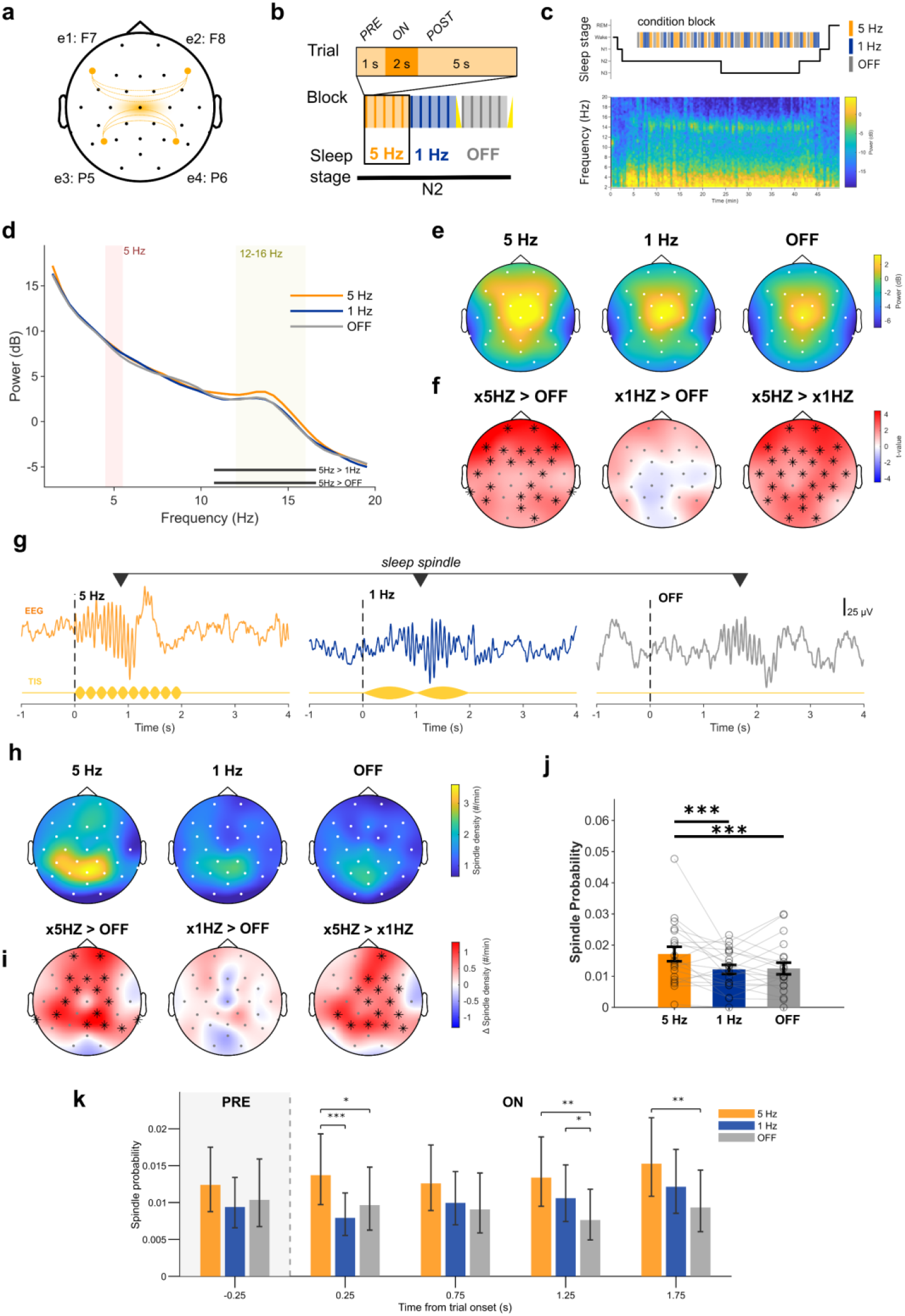
Experimental results (a) TIS electrode montage: Pairwise connected electrodes were attached at scalp positions F7 (e1) and F8 (e2), and at positions P5 (e3) and P6 (e4). (b) Stimulation protocol with 1Hz, 5Hz, and OFF conditions applied during N2/N3 sleep. Stimulation periods are 2s long with a 6s refractory period. Stimulation frequencies were df =5 Hz with cf = 2000 Hz, df = 1 Hz; with cf = 2000 Hz, at 2 mA peak-to-baseline, and df=0Hz with cf= 0 Hz at 0 mA (OFF). (c) Example stimulation schedule in one participant. Stimulation blocks (top are shown relative to the hypnogram and example multitaper spectrogram (channel C3). (d) Average multitaper power spectrum across all electrodes. Cluster corrected mass-univariate testing shows a significantly higher sigma power (10.7 – 16.6 Hz) in the 5Hz condition compared to 1Hz and OFF condition (black bars, Table S5, S6) but no power difference in theta frequency (4-6 Hz) range (see also theta power topography Table S9, S10). (e) Topographical plots of electrode-wise fast spindle power (FSP, 12-16 Hz) derived from multitaper spectra. (f) Topographical differences in FSP revealed a nearly uniform increase in FSP in the 5Hz condition (cluster corrected one sample t-test,5Hz > 1Hz, cluster mass = 64.83, 26/30 electrodes, p = 0.009; 5Hz > OFF: cluster mass = 55.85, 23/30 electrodes, p = 0.018, Table S7, S8). (g) Example raw EEG traces of spindle events detected with the YASA toolbox for each experimental condition. TIS trace represented below as an envelope amplitude waveform that is modulated periodically at df.(h) Topographical plots of fast spindle density showing canonical centro-pariatal distributions s. (i) Topographical difference plots revealed an increase in spindle density in the 5Hz condition at central/parietal electrodes (cluster corrected one sample t-test, 5Hz > OFF: cluster mass = 33.12, 15/30 electrodes, p = 0.018, 5Hz > 1Hz, cluster mass = 30.63, 16/30 electrodes, p = 0.022, Table S15, S16). (j) Spindle probability analysis: A generalized linear mixed model employing a binomial modelwith logit link, fitted with maximum likelihood using Laplace approximation. The binomial model results confirm a main effect of condition (Condition: ChiSq: 40.64, df=2, p<0.001), with significantly higher spindle probability in the 5Hz compared to 1Hz and OFF conditions after post-hoc pairwise comparison with FDR correction (5 Hz>OFF, OR: 0.84, SE:0.03,z=-5.24, p<0.001, 5Hz >1 Hz OR: 0.84, SE: 0.03z= 5.54,p<0.001, Table S17, S18). (k) Post stimulus histogram of spindle probability: spindle event probabilities in 500 ms bins during trials. Higher probability for spindle events occurs in the 5Hz condition compared to 1 Hz and OFF conditions immediately after stimulus onset (0-0.5 s post stimulus onset, bin center: 0.25: contrast:5 Hz>1 Hz, OR:0.58, SE: 0.08, z=-4.11, p< .001; contrast 5 Hz>OFF, OR:0.70, SE: 0.11, z=-2.19, p=0.043)) and in windows 1-1.5 s post stimulus onset (Table S19, S20, S21).

Multitaper spectral analysis revealed a significant increase in spindle-band (sigma) power during the 5Hz condition compared to the 1Hz and OFF conditions (Figure 2d, subject-level cluster-corrected one-sample t-statistic on within-subject differences, 5Hz>1Hz: cluster mass=29.42, 10.74-16.60Hz; p=0.010, 5Hz > OFF: Cluster mass=29.42, 10.74-16.60Hz, p=0.017, 1Hz>OFF: n.s., Table S5, 6).

We focused the remainder of our analysis on fast spindle power and fast spindle activity based on intrinsic frequency (12-16 Hz)^1^.

Fast spindle power (FSP, 12-16 Hz) topographies showed typical centro-parietal distributions (Figure 2e), with nearly uniform increase in FSP in the 5Hz condition electrodes (cluster corrected one sample t-test, 5Hz > 1Hz, cluster mass=64.83, 26/30 electrodes, p=0.009; 5Hz > OFF: cluster mass=55.85, 23/30 electrodes, p=0.018, Figure 2f). Stimulation-onset spectrograms confirmed enhancement was spectrally specific to the fast sigma band and temporally aligned with stimulation epochs, supporting a selective modulation spindle activity rather than a broadband increase in EEG power (Figure S4e).

To determine whether the observed increase in FSP was driven by augmented spindle generation, spindle events were automatically detected using the YASA toolbox^20^. Fast spindle events did not differ in morphology or in derived properties such as amplitude, frequency or duration across conditions (Figure S5a, Table S13, S14).

Fast spindle density (FSD, events/min) was higher during 5Hz (linear mixed model, F (2,1797), 3.12, p=0.044, Figure S5b, Table S15) compared to 1Hz and OFF condition. Topographical plots showed a typical centro-parietal distribution of FSD (Figure 2h), which was increased in the 5Hz condition at central/parietal electrodes (cluster corrected one sample t-test, 5Hz > 1Hz, cluster mass=64.83, 26/30 electrodes, p=0.009; 5Hz > OFF: cluster mass=55.85, 23/30 electrodes, p=0.018, Figure 2i, Tables S18-19).

A binomial mixed model using binary spindle outcome per trial confirmed the main effect of condition (ChiSq: 40.64, df=2, p<0.001), with significantly higher spindle probability in the 5Hz compared to 1Hz and OFF conditions (5Hz>OFF, OR: 0.84, SE:0.03,z=-5.24, p<0.001, FDR corrected, Figure 2j, Table S18). A post stimulus time histogram of spindle event probability computed over 500ms windows (Figure 2k) showed a higher probability for spindle events in the 5Hz condition immediately after stimulus onset (0-0.5s post stimulus onset, bin centre: 0.25: contrast:5Hz>1Hz, OR:0.58, SE: 0.08, z=-4.11, p< .001; contrast 5Hz>OFF, OR:0.70, SE: 0.11, z=-2.19, p=0.043, FDR corrected, Table S26) and in windows 1-1.5s post stimulus onset (Table S21).

To delineate the mechanistic origin of TIS frequency selectivity, we examined a biophysical model of an isolated TC neuron. With minimal external stimulation, the model exhibited transient subthreshold oscillations near 5Hz across physiologically realistic T-type calcium and h-current conductances (Figure 3a).

**Figure 3.**
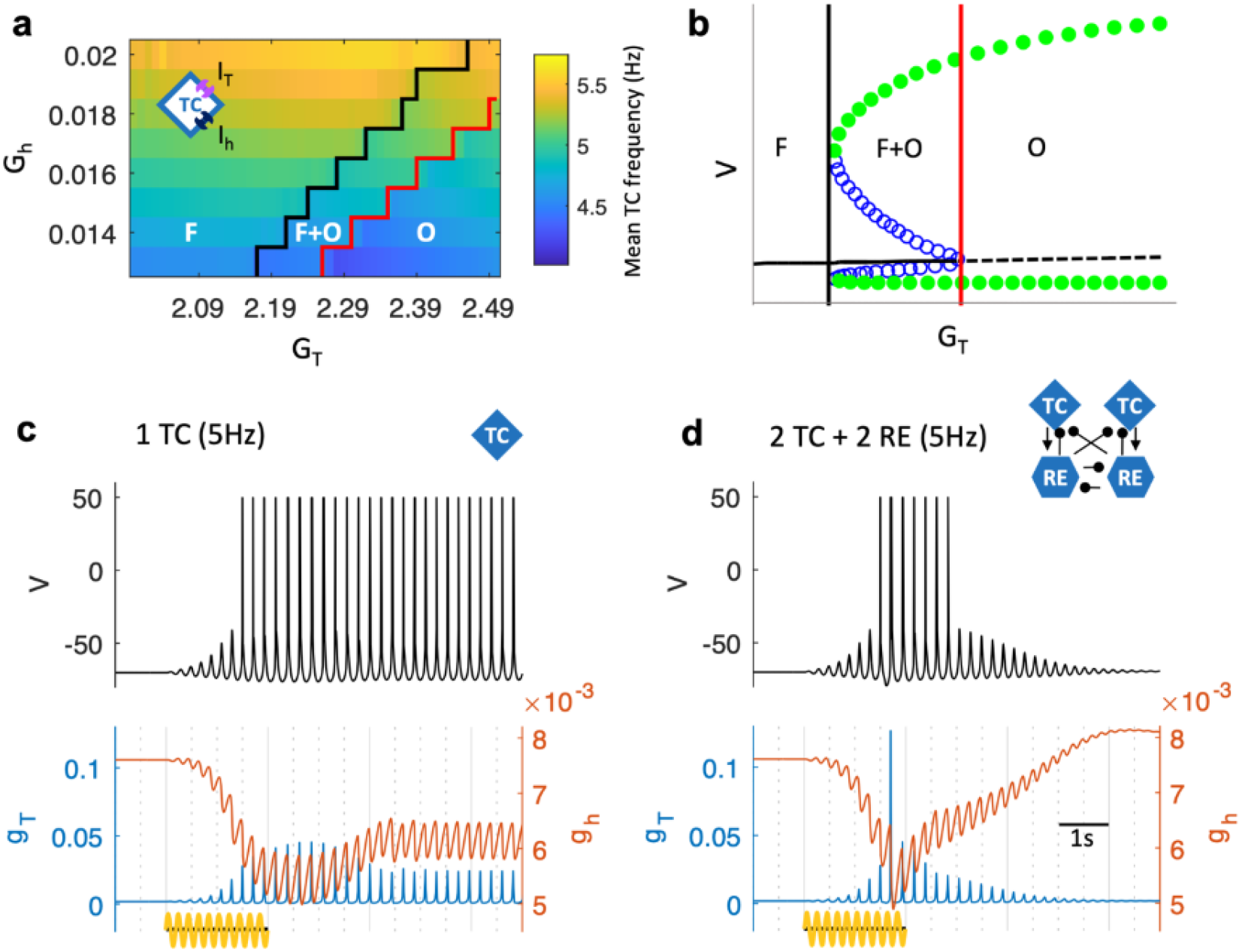
Minimal computational model results (a) Native oscillation frequency of isolated TC model cells for a range of T- and h-current maximum conductance values (G_T_ and G_h_, respectively). Membrane potential dynamical regimes are separated by solid lines and labeled as fixed point (“F”) at resting potential, coexistence of stable fixed point and sustained spiking oscillations (“F+O”), and oscillations only (“O”). (b) Bifurcation of membrane dynamics with respect to T-conductance G_T_ for G_h_=0.016 in a simplified isolated TC model (see Methods). Fixed-point (“F”), bistability (“F+O”), and stable limit cycle (“O”) regimes are delimited with vertical solid lines. (c) 5Hz 2s sub-threshold stimulation (sine wave, 0.01 *μ*A intensity) for an isolated TC cell in the bistable regime (G_T_=2.34, G_h_=0.016). From top to bottom: membrane voltage, calcium influx, T current and h current. Sub-threshold oscillations are amplified into the spiking regime. (d) As (c), for one TC cell in a minimal circuit with 2 RE and 2 TC interconnected cells. The resonance near 5Hz holds, but RE bursts (not shown) induce a calcium peak that up-regulates h-current effective conductance (gh) and terminates the spindle.

To understand why subthreshold stimulation can trigger suprathreshold oscillations, we performed bifurcation analysis of a reduced model in which h-current conductance is independent of Ca^2+^ concentration (Figure 3b). This revealed bistability between resting (stable fixed point, F) and oscillatory (limit cycle, O) states across a range of T-current conductances (F+O region, Figure 3b). Within this bistable regime, sinusoidal input near the resonant frequency induced subthreshold membrane oscillations, which were then amplified until spiking occurred, leading to sustained supra-threshold oscillations in the spindle frequency range (Figure3c). Inputs of identical amplitude at non-resonant frequencies decayed to rest without amplification (Figure S6).

In a minimal circuit model (2 TC + 2 RE neurons, Figure 3d), TC neuron resonance governed spindle initiation, while calcium influx via the T-channels, upregulating h-current conductance, drove spindle termination^13,14^. Importantly, bistability was maintained so once oscillations became suprathreshold, they triggered a full spindle event. Thus, TC subthreshold resonance acts as a nonlinear amplifier that selectively converts subthreshold oscillatory input into spiking, triggering a spindle.

The model predictions align with prior evidence linking TC resonance^21^ to circuit oscillations, including subthreshold membrane oscillations^22^ and slow oscillatory firing modes^23^. Previous studies showed that rhythmic suprathreshold stimulation can evoke augmented thalamic responses, dependent on T-channel-mediated low-threshold Ca^2+^ spikes, when stimulation frequency matches intrinsic rebound timescales^24^. Here, we extend this framework to the subthreshold regime, showing that the interaction of T- and h-currents defines a preferred frequency range for TC responses to external electrical fields, while network bistability can enable weak stimulation to trigger large-amplitude spindle events, increasing spindle density.

Our experimental results confirm a frequency specific rather than non-specific effect of non-invasive thalamic TIS, whereby 5Hz was most effective compared to a 1Hz active control condition. The estimated field strengths of approximately 0.4 V/m have previously been associated with subthreshold modulation of membrane potential and spike timing^25^.

Several limitations should be acknowledged. First, the computational model necessarily simplifies the full complexity of the human thalamocortical system; though its core mechanisms for T- and h-currents, universally expressed in thalamic nuclei^26^, likely produce dynamics that are robust to parameter variations and scales. Second, we only provide indirect evidence of thalamic engagement, without individualized FEM modelling or invasive thalamic recordings. Finally, a limited set of envelope frequencies was tested experimentally, constraining conclusions about the full frequency-response profile. The model predicts a second resonance regime near 10Hz, consistent with recent findings that thalamic TIS at this frequency can also modulate spindle activity^27^.

Together, our findings support the idea that intrinsic biophysical properties of thalamocortical neurons can be exploited for selective, non-invasive modulation of sleep spindles. By linking stimulation parameters to candidate cellular resonance mechanisms, this work provides a mechanistic basis for refining stimulation protocols and developing interventions aimed at enhancing sleep-dependent memory processing and ameliorating spindle deficits in clinical populations.

## Methods

### Computational Modelling

#### Network Geometry

The thalamocortical network model followed the structure published in Wei et al.^28^ with minor modifications. It included 1,000 pyramidal (PY), 200 inhibitory (IN), 500 reticular (RE), and 500 thalamic relay (TC) neurons with individual dynamics based on the Hodgkin-Huxley kinetics^29^. Each population was organized in a one-dimensional layer projecting to other layers through local synaptic connections. PY neurons projected to both PY and IN cells with AMPA and NMDA connections, while IN neurons projected to PY cells only through GABA_A_ connections. Within the thalamus, RE cells received AMPA connections from TC neurons as well as GABA_A_ synapses from other RE cells, whereas TC cells had only inhibitory GABA_A_ and GABA_B_ inputs from RE cells. Thalamocortical connectivity consisted of AMPA synapses from TC to PY and IN neurons, as well as from PY to TC and RE cells. Connection radii are shown in Table 1 and are identical to those in Wei et al.^28^.

**Table 1.** Connection radii between different populations in the thalamocortical model, corresponding to the difference between neuron linear indices.

| From | To | Synaptic type | Radius |
| --- | --- | --- | --- |
| PY | PY | AMPA | 5 |
| PY | PY | NMDA | 5 |
| PY | IN | AMPA | 1 |
| PY | IN | NMDA | 1 |
| IN | PY | GABA<sub>A</sub> | 5 |
| TC | RE | AMPA | 8 |
| RE | TC | GABA<sub>A</sub> | 8 |
| RE | TC | GABA<sub>B</sub> | 8 |
| RE | RE | GABA<sub>A</sub> | 5 |
| TC | PY | AMPA | 15 |
| TC | IN | AMPA | 3 |
| PY | TC | AMPA | 10 |
| PY | RE | AMPA | 8 |

#### Intrinsic currents: Cortex

The cortical neurons included dendritic and axo-somatic compartments.

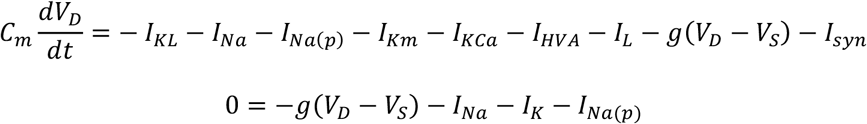

where *C*_*m*_ is the membrane capacitance, *I*_*KL*_ is potassium leak current, *I*_*Na*_ is a fast sodium current, *I*_*Na*(*p*)_ is a persistent sodium current, *I*_*Km*_ is a slow voltage-dependent non-inactivating potassium current, *I*_*KCa*_ is a slow Ca^2+^-dependent K+ current, *I*_*HVA*_ is a high-threshold Ca^2+^ current, *I*_*L*_ is the Cl-leak current, *g* is the conductance between axo-somatic and dendritic compartment. *V*_*D*_ and *V*_*S*_ are the membrane potentials of dendritic and axosomatic compartments, and *I*_*syn*_ is the sum of synaptic currents to the neuron.

The persistent sodium current *I*_*Na*(*p*)_ was included in the axosomatic and dendritic compartment of PY cells to increase bursting propensity. IN cells had the same intrinsic currents as those in PY except that *I*_*Na*(*p*)_ was not included. All the voltage-dependent ionic currents *I*_*j*_ have the canonical form:

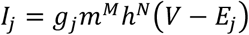

where *g*_*j*_ is the maximal conductance, *m* and h are dynamic gating variables, *V* is the voltage of the corresponding compartment and *E*_*j*_ is the reversal potential. See Wei et al.^28^ for further details.

#### Intrinsic currents: Thalamus

TC and RE cells were modeled as single compartments with membrane potential *V* governed by the Hodgkin-Huxley kinetics, so that the total membrane current per unit area is given by

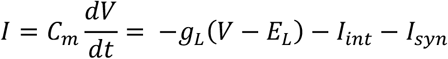

Where *C*_*m*_ is the membrane capacitance, *g*_*L*_ is the non-specific (mixed Na^+^ and Cl^-^) leakage conductance, *E*_*L*_ is the reversal potential, *I*_*int*_ is a sum of active intrinsic currents, and *I*_*syn*_ is a sum of synaptic currents.

Intrinsic currents for both RE and TC cells included a fast sodium current, *I*_*Na*_, a fast potassium current, *I*_*K*_, a low-threshold Ca^2+^ current, *I*_*T*_, and a potassium leak current, *I*_*KL*_ = *G*_*KL*_(*V* − *E*_*K*_), where *G*_*KL*_ is the maximum potassium leakage conductance and *E*_*K*_ is the potassium reversal potential. For TC cells, an additional potassium A current, *I*_*A*_, and a hyperpolarization-activated cation current, *I*_*h*_, were also included. All voltage-dependent ionic currents *I*_*j*_ (*j* = *Na, K, T, A*) have the general form

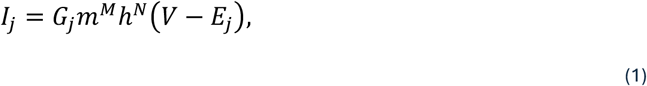

where *g*_*j*_ is the maximal conductance, *E*_*j*_ is the reversal potential, and *m* and *h* are dynamic gating variables. The detailed dynamics of gating variables and their associated transition rates are described in Bazhenov et al.^24^.

##### Low-threshold calcium T-current

The T current follows the general form in Eq. (1) for intrinsic currents, with calcium-dependent reversal potential based on the Nernst equation:

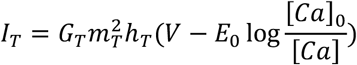

where [*Ca*] is the calcium concentration, and *E*_0_ = *RT*/2*F* with *R* =8.31441J/(mol K), *T* = 309.15 K, *F* = 96489 C/mol, and [*Ca*]_0_ is the extracellular calcium concentration.

##### Calcium concentration

The calcium dynamics are given by

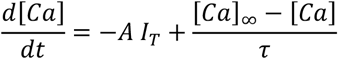

where [*Ca*]_∞_ is the equilibrium calcium concentration, A is a calcium drive modulation constant and *τ* is the time constant for calcium depletion.

##### Hyperpolarization-activated cation current

The h current *I*_*h*_ incorporates both calcium and voltage dependencies. *I*_*h*_ is described by

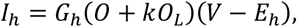

where *G*_*h*_ is the maximal conductance, *E*_*h*_ = −40 mV is the reversal potential, k=2 is a constant, and *O* and *O*_*L*_ are the fraction of open and locked *h* channels, respectively.

The voltage dependence is reflected in the first-order kinetics of the transition between closed *C*and open *O* states of h-channels without inactivation

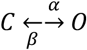

where *α*(*V*) and *β*(*V*) are voltage-dependent transition rates. Calcium dependence is implemented in the higher-order kinetics governing the transition to the locked form *O*_*L*_, which involves an intermediate regulation factor *P*. The binding of calcium ions with the unbound form of the factor, *P*_0_, leads to the bound form *P*_1_, which binds to open channels *O* to produce the locked form *O*_*L*_.

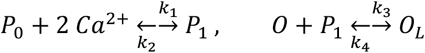

where *k*_1_, *k*_2_, *k*_3_, *k*_4_ are constant rates.

Detailed expressions for all currents are given in Bazhenov et al.^30^. The main model parameters in Table 2 are largely identical to those in Wei et al.^28^, where they were determined based on previous modeling studies grounded in experimental observations^30,31^. Specifically, thalamic reticular and relay neurons included intrinsic properties necessary for generating rebound responses which were found to be critical for spindle generation.

**Table 2.** Parameter values for thalamic cells in the thalamocortical model.

| Parameter | Description | TC value | RE value | Units |
| --- | --- | --- | --- | --- |
| $S$ | Cell surface | $2.9 \cdot 10^{-4}$ | $1.43 \cdot 10^{-4}$ | $\text{cm}^2$ |
| $C_m$ | Membrane capacitance | 1 | 1 | $\mu\text{F}/\text{cm}^2$ |
| $G_L$ | Leak maximal conductance | 0.01 | 0.05 | $\text{mS}/\text{cm}^2$ |
| $E_L$ | Leak reversal potential | -70 | -77 | mV |
| $G_{KL}$ | $\text{K}^+$ leak maximal conductance | 0.023 | 0.012 | $\text{mS}/\text{cm}^2$ |
| $E_K$ | $\text{K}^+$ reversal potential | -95 | -95 | mV |
| $G_T$ | T-current maximal conductance | 2.4 | 2.2 | $\text{mS}/\text{cm}^2$ |
| $[Ca]_0$ | Extracellular $\text{Ca}^{2+}$ concentration | $10^{-4}$ | $10^{-4}$ | mM |
| $G_h$ | h-current maximal conductance | 0.016 | - | $\text{mS}/\text{cm}^2$ |
| $E_h$ | h- current reversal potential | -40 | - | mV |
| $[Ca]_\infty$ | $\text{Ca}^{2+}$ equilibrium concentration | $2.4 \cdot 10^{-4}$ | $2.4 \cdot 10^{-4}$ | mM |
| $A$ | $\text{Ca}^{2+}$ drive modulation | $2.5 \cdot 10^{-5}$ | $5 \cdot 10^{-5}$ | $\text{mM cm}^2/(\text{ms } \mu\text{A})$ |
| $\tau$ | $\text{Ca}^{2+}$ depletion time | 5 | 5 | ms |

#### Synaptic currents

Synaptic currents (GABA, NMDA and AMPA) were modeled by first-order activation schemes, using the general form

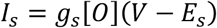

where *s* is AMPA, NMDA, GABA_A_ or GABA_B_, *g*_*s*_ is the maximal synaptic conductance, [*O*] is the time-dependent fraction of open channels, and *E*_*s*_ is the reversal potential. See Wei et al.^28^ for details.

#### Simulated stimulation

The effect of temporal-interference stimulation (TIS) was simulated as a sinusoidal input current *I*_*TIS*_ at the TIS envelope frequency. For thalamic stimulation, all TC and RE model cells received inputs. The amplitude of the injected sinusoidal current was maximal at the layer midpoint and decayed slowly with distance as a gaussian, so that for a cell with index *i*,

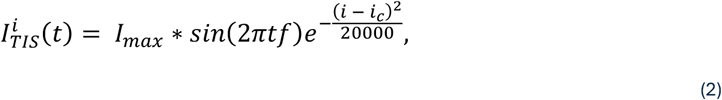

where *I*_*max*_ is the maximal stimulus intensity (sine wave amplitude), *f* is the stimulus frequency, *i* is the cell index and *i*_*c*_ is the index of the target cell, which in this case is the center of the layer (250 for the 500-cell thalamic layers).

We conducted a systematic sweep over stimulus frequencies and intensities to identify the combinations with best spindle responses in the model. To isolate the effect of stimulation on thalamic spindle response, we removed all cortical inputs to the thalamus by setting the synaptic strengths of all PY-TC and PY-RE connections to 0. Thalamocortical projections were left intact, and we recorded spindle responses from the cortex. The stimulation frequency *f* was varied from 1Hz to 15Hz, and the range of maximal intensities *I*_*max*_ was 0.01 to 0.05 *μ* A. All intensities were subthreshold, meaning that the injected current magnitude was not large enough to immediately push the membrane potential over the spiking threshold. The voltage deviation from resting potential directly induced by the maximum current injected (0.05 *μ* A) was under 2mV. Simulations for each combination were run for 40s, with a single 2s stimulation period at time 10s.

To compare the effect of stimulation frequency on spindle density, simulation duration was set to 610s and 1Hz or 5Hz stimulation was applied for 2s, every 8s (6s refractory period), with *I*_*max*_ = 0.03 *μ*A. Each condition (1Hz, 5Hz, or no stimulation) was replicated 9 times using different random seeds. This randomness affects intrinsic cell properties and initial conditions, but not connectivity.

We compared the model’s response to stimulation when targeting different cell populations. For this, we used simulations 610s long with 1Hz or 5Hz stimulation applied for 2s every 8s with *I*_*max*_ = 0.03 *μ* A. Stimulation intensity followed the same gaussian field in Eq. (2) centered at the middle cell of the target layer or layers. In addition to thalamic stimulation targeting all TC and RE cells, we ran simulations targeting each population separately (only TC, RE, PY, or IN), as well as PY and IN together.

#### Detection of model spindles

Spindle detection for the model generally followed the methodology in Purcell et al. ^32^ and used custom code implementing Morlet wavelet analysis in MATLAB. To account for variance in spindle frequency, a filter bank of Morlet wavelets with frequencies in the 9-16Hz range was created. Each LFP channel, which represents the average voltage of 100 consecutive cells, was filtered in the 9-16Hz band and detrended. For each channel, the continuous wavelet transform (CWT) was obtained using the filter bank. The wavelet coefficients were averaged over frequencies. The average coefficients were smoothed over time using a 100ms-window moving average and normalized between 0 and 1. Points where this wavelet signal exceeded a multiplicative threshold of 3 times the baseline, defined as the median of the signal, were flagged. Intervals of consecutive flagged points with durations greater than 0.3s were flagged as putative spindle cores, and cores closer than 0.5s were merged. To capture the characteristic waxing-and-waning profile of spindles, we extended each core with flanks of consecutive surrounding points exceeding 1.5 times the baseline. Extended cores were discarded if their duration was under 0.5s, and merged if their separation was less than 0.5s. The resulting events were considered detected spindles.

#### Properties of model spindles

Local activity channels were obtained by averaging the voltage traces of groups of 100 consecutive PY neurons (10 groups). Spindles were detected on filtered signals from each channel. For each spindle detected, amplitude was set to the maximum peak-to-peak difference, and the frequency was estimated as the inverse of the within-spindle inter-peak time in seconds.

For each channel, density was estimated as the total count of spindles over the total simulation time in minutes. Density was averaged over channels for each simulation. To compare density across stimulation conditions in Figure 1h, we first tested for normality within conditions using an Anderson-Darling test. Because the density distributions were not normal, we conducted a Wilcoxon rank-sum test on each pair of conditions, with each condition having 9 simulations.

#### Phase-locking to model stimulation

To determine how strongly the timing of spindles entrained to the timing of stimulation, we computed the phase of each spindle onset with respect to the 8s period between stimulus intervals. The individual phase *ϕ*_*i*_ for spindle *i* was obtained as

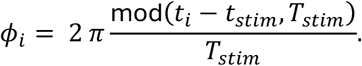

where *t*_*i*_ is the spindle onset, *t*_*stim*_ is the time of the first stimulation interval, and *T*_*stim*_ = 8s is the period between stimulation intervals. The phase locking value was obtained for each channel as

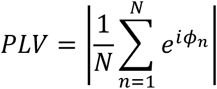

where N is the number of spindles detected in the channel.

#### Isolated thalamocortical (TC) neuron model

To characterize the intrinsic oscillatory and resonance properties of thalamocortical (TC) relay neurons, we performed simulations of a single isolated TC model neuron. Each simulation lasted 100s and included a single perturbation applied at 20s, allowing the membrane potential to fully relax to its resting state before stimulation. Perturbations consisted of either a brief current pulse or a sinusoidal input, depending on the analysis, and were designed to probe the neuron’s subthreshold and spiking dynamics.

For the analysis shown in Figure 3a, we applied a 500ms current pulse with amplitudes ranging from 0.001 to 0.1 *μ*A, which included both sub- and super-threshold perturbations. Varying the perturbation amplitude allowed us to distinguish between dynamical regimes in which the neuron returned to rest after perturbation and those in which sustained oscillatory activity emerged. The dominant oscillation frequency for each simulation was defined as the frequency of the largest peak in the power spectrum of the detrended membrane potential trace. Power spectra were computed using MATLAB’s pspectrum() function over the 0.2 - 20Hz range. All simulations were sampled at 1000Hz.

Simulations were repeated across combinations of T-type calcium conductance (*G*_*T*_, 0.013– 0.020 mS/cm^2^) and h-current conductance (*G*_*h*_ = 2.0 - 2.5 mS/cm^2^). For each (*G*_*h*_, *G*_*T*_) pair, the mean intrinsic TC frequency reported in Figure 3a corresponds to an average across all perturbation amplitudes tested. This averaging captures the characteristic subthreshold timescale of the neuron even in regimes where oscillations are not self-sustaining.

Simulations with different perturbations were run for combinations of T-conductance *G*_*T*_ between 2.0 and 2.5 mS/cm^2^ and h-conductance *G*_*h*_ between 0.013 and 0.02 mS/cm^2^. The mean TC frequency reported for each combination is an average over perturbation amplitudes.

The black and red lines in Figure 3a indicate empirically determined boundaries between dynamical regimes based on the neuron’s state at the end of each 100s simulation. Simulations to the left of the black curve (region “F”) always converged to a single stable fixed point corresponding to the resting membrane potential. Between the black and red lines (region “F+O”) trajectories either converged to resting potential or exhibited oscillations depending on the perturbation amplitude. Simulations to the right of the red curve (region “O”) failed to return to rest within 100s and instead developed oscillations.

The bifurcation diagram shown in Figure 3b illustrates the origin of these regimes using a reduced version of the TC model. To isolate subthreshold dynamics, we removed fast sodium and delayed rectifier potassium currents responsible for action potential generation. To obtain stable sustained oscillations rather than waxing-and-waning activity, we additionally removed the calcium dependence of the h-current. Finally, we omitted the A-type potassium current, which has relatively small conductance and does not play a primary role in spindle dynamics. The resulting simplified system captures the essential interaction between T-type calcium and h-currents underlying TC resonance.

Bifurcation analysis was performed using XPPAUT, with *G*_*T*_ as the bifurcation parameter and *G*_*h*_ fixed at 0.016. Vertical lines in Figure 3b mark the locations of a subcritical Andronov–Hopf bifurcation (red) and a fold of limit cycles (black). Across the parameter range explored in Figure 3a, changes in *G*_*h*_ shifted the locations of these bifurcation points but did not alter the qualitative structure of the diagram. The full TC model exhibits the same sequence of dynamical regimes as *G*_*T*_ increases (stable fixed point, coexistence of fixed point and oscillations, and oscillations only) although the oscillations are more complex due to the calcium dependency of the h-current and are beyond the scope of the present analysis. Here, we focus specifically on the resonance properties that allow subthreshold inputs to drive the system out of the resting fixed-point basin and toward spiking. In a network context, the first TC spike recruits additional cells and reshapes the oscillation through synaptic interactions.

The region where the resting attractor and stable oscillations coexist (“F+O” in Figure 3a) is the regime in which stimulation properties can determine the system’s response, and thus where frequency selectivity is expected to emerge. To demonstrate this, we repeated isolated TC simulations for *G*_*h*_=0.016 and *G*_*T*_=2.34, replacing the 500ms pulse perturbation with a 2s sinusoidal input of identical amplitude (0.01 *μ*A) delivered either at 5Hz or 1Hz. These simulations correspond to Figure 3c and Figure S6a, respectively.

The minimal circuit model shown in Figure 3d consisted of two TC and two RE neurons connected via RE–RE and RE–TC synapses with the same properties as in the full thalamocortical network. The simulation protocol was identical to that used for the isolated TC neuron in Figure 3c, with 2s sub-threshold sinusoidal inputs at either 5Hz (in Figure 3d) or 1Hz (Figure S6b) applied to all TC and RE neurons. This minimal circuit was used to confirm that the resonance-driven initiation mechanism observed in isolated TC neurons persists in the presence of reciprocal TC–RE interactions.

### Experimental Methods

#### Participants

We recruited 25 right-handed adults from within the University of Surrey via posters, e-mails, and the university student recruitment system SONA. Five participants did not achieve stable N2 or N3 sleep during the nap session and were excluded from all analyses, yielding a final sample of 20 participants (mean age=27 years; range 19-35 years).

Inclusion criteria required participants to be aged 18 years or older with a body mass index below 30. Exclusion criteria comprised any history of neurological, psychiatric, or sleep disorders; current medical illness requiring medication; substance abuse or alcohol consumption exceeding 28 units per week within the previous six months; personal or family history of epilepsy; history of migraine; unexplained loss of consciousness or brain injury; prior neurosurgery; implanted electronic or intracranial metal devices; scalp skin conditions, burns, or injuries; night-shift work or travel across >2 time zones within the preceding month; and pregnancy. All participants were registered with a general practitioner and provided written informed consent prior to participation.

The study was approved by the University of Surrey Ethics Committee (FHMS 22–23 123 EGA / ERM 873). Participants received £40 for the nap visit and reimbursement of travel expenses up to £20.

#### Procedure

Each participant attended a single daytime session at the Surrey Sleep Research Centre, beginning at 2 pm. Participants were instructed to maintain regular sleep schedules in the days preceding the visit, to avoid alcohol for 24 hours prior to testing, and to abstain from caffeine on the day of the session. On arrival, participants were asked to attempt to sleep as they usually would during a daytime nap.

Upon arrival, stimulation electrode sites were attached to the head, and stimulation intensity was individually titrated (see below, *Temporal Interference Stimulation*). Then EEG, EMG and EOG electrodes were applied. Participants asked to sleep in a darkened, sound-insulated, temperature-controlled sleep laboratory pod at approximately 3 pm. The nap opportunity lasted up to 90 minutes (mean=75 ± 11 minutes). The experimenter monitored the polysomnographic recording in real time from an adjacent control room and initiated the stimulation protocol once stable N2 or N3 sleep was established (see *Experimental Design*). Stimulation was paused upon any arousal and resumed when stable sleep was re-established. The session was terminated when the 90-minute maximum was reached, or at the experimenter’s discretion once the participant was likely to no longer sleep.

#### Experimental Design

A within-subject design was employed in which each participant received all three stimulation conditions during a single nap session: (i) TIS with a 5Hz envelope (f1=2000Hz, f2=2005Hz, df=5Hz, 2mA per channel pair), (ii) TIS with a 1Hz envelope (f1=2000Hz, f2=2001Hz, df= 1Hz, 2mA per channel pair), and (iii) sham stimulation (OFF; 0Hz, 0 mA).

The 5Hz condition was the active condition of primary interest, designed to target subthreshold resonance in corticothalamic relay neurons. The 1Hz condition served as active control, with no predicted subthreshold resonance. The OFF condition served as control condition.

##### Trial and block structure

Each trial lasted 8-seconds (Figure 2b): An ON trial consisted of 1 second PRE window (f1=f2=2000Hz, df=0), 2 seconds of envelope-modulated temporal interference stimulation (df =5 or 1Hz) followed by 5 seconds POST window (f1=f2=2000Hz, df=0). An OFF trial consisted of 8 seconds with no current applied (f1=f2=0Hz, df=0).

Five consecutive trials of the same condition formed one condition block (total duration =40 seconds). Block order was counterbalanced across participants using a Latin-square design. For each participant, a pre-generated block sequence file specified the ordering of blocks. Three stimulation conditions (5Hz, 1Hz, OFF) appeared in a pseudorandomised order, with each condition presented for five consecutive trials before switching to the next condition. Sequences of three blocks were repeated for the duration of the nap. This design minimised systematic order and carry-over effects across participants. When a change from OFF to ON or ON to OFF blocks, a 5 second ramp up or ramp down window was inserted between blocks.

##### Stimulation protocol

Stimulation was initiated manually by the experimenter based on real-time polysomnographic monitoring. The experimenter confirmed stable NREM sleep, defined as continuous N2 or N3 staging for at least four consecutive 30-second epochs (≥ 2 minutes) before starting the stimulation protocol. If the participant showed signs of arousal or transitioned to N1, Wake, or REM, stimulation was immediately paused. Stimulation was resumed once stable N2 or N3 sleep was re-established using the same criterion.

The number of stimulation trials per condition varied across participants depending on total time spent in N2/N3 sleep (mean=95 ± 35 trials per condition).

#### Blinding

##### Participant blinding

Prior to the nap, stimulation intensity was individually titrated to ensure imperceptibility (see *Titration*). Following titration and scalp preparation, all participants confirmed that they could not perceive stimulation at the experimental intensity of 2 mA.

##### Experimenter and analysis blinding

Sleep scoring was performed blind to stimulation condition and trial timing. The hardware low-pass filter applied during recording (see *EEG Recording*) attenuated high-frequency carrier components, preventing the scorer from identifying active stimulation epochs in the EEG trace. Only ramping artefacts at condition transitions were visible, but these did not reveal which condition preceded or followed the ramp. All primary analyses employed fully automated, pre-specified pipelines for EEG preprocessing, spindle detection, trial assignment, and statistical modelling.

#### Temporal Interference Stimulation and EEG

##### Field modelling

Electromagnetic (EM) simulations were performed using the Temporal Interference Planning tool (https://tip.science, IT’IS Foundation, Zurich, Switzerland) and Sim4Life (ZMT Zurich MedTech AG, Zurich, Switzerland).

All calculations were performed using the structured (rectilinear) version of the Low-Frequency Quasi-Static Ohmic Current Dominated (LF-QS-OCD) solver from Sim4life.

The LF-QS-OCD solver finds the solution to the QS-approximation of Maxwell’s equations:

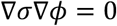

where σ is the local electrical conductivity (scalar or tensorial) and ϕ is the electric potential (electric field: E=-∇ϕ).

The QS-approximation is valid when:

> σ >> ωϵϵ_0_, where ϵ is the electrical permittivity of the tissue/material, ϵ0 is the permittivity of the vacuum, and ω is the angular frequency of the applied field, and
>
> λ >> d, where λ is the wavelength of the EM field and d is the characteristic size of the structure of interest or the computational domain.

Under these conditions, ohmic currents dominate over displacement currents, and inductive effects can be neglected.

The final electrode montage was derived using TIP (using a Sim4Life backend) with the MIDA head model ^33^ with additional segmentation of deep brain regions.

TIP uses a multigoal optimisation to optimise the predicted targeting of the bilateral whole-thalamus as target region.

We used the modulation envelope magnitude 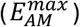 as TI-exposure quantity, which is computed according to the formula by Grossman et al.^18^:

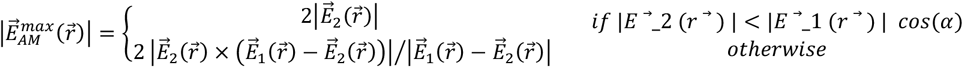

To assess the quality of a TI exposure condition, three key metrics are evaluated simultaneously: 1) Target exposure strength: the 50th isopercentile of 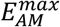 in the target, 2) Exposure selectivity: the ratio of the mean target 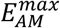 and the mean off-target 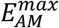, 3) Off-target exposure (collateral): the fraction of the non-target brain volume with 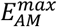 exceeding target exposure strength.

These metrics are leveraged by a Pareto multi-goal optimization algorithm to determine Pareto-optimal electrode positions that minimise all 3 metrics simultaneously. Here ‘Pareoto-optimal’ means that no configuration can be found that improves on one or multiple quality metrics, without worsening performance on at least one other metric.

After optimisation, all electrode pair combinations are ranked according to dynamically adjustable weights for the different metrics. From the evaluation of several million possible stimulation conditions, we chose a montage that maximises exposure strength with reduced selectivity and increased collateral metrics (Figure S3b).

##### Device and safety

Stimulation was delivered using a galvanically isolated, eight-channel high-precision current-source stimulator (TIBS-R V3.0, TI Solutions AG, Switzerland). The channels are fully synchronized, as they are synthesized by the same field-programmable gate array (FPGA), and output currents as well as electrode impedances are continuously monitored.

Fully characterized hardware filters were used to attenuate stimulation artifacts in the EEG to below detectable levels (<30 nV RMS). An 8-channel passive current-mode high-pass filter (HPF-8, IT’IS Foundation, Switzerland) was connected between the stimulator output and the stimulation electrodes. In addition, a 7th-order low-pass filter (LPF-32BP, IT’IS Foundation, Switzerland), with a cutoff frequency of 90 Hz and >130 dB attenuation above 1 kHz, was inserted between the EEG electrodes and the EEG amplifier. This set up minimises intermodulation artefacts at the difference frequency in EEG recordings.

##### Stimulation parameters

The carrier frequency was fixed at 2000 Hz on both channel pairs, with the second pair offset by the desired envelope frequency (2005 Hz for the 5Hz condition, 2001 Hz for the 1Hz condition). Current amplitude was 2mA (zero-to-peak) per channel pair for all active conditions. In the OFF condition, both channels were set to 0Hz and 0 mA.

Transitions between conditions involving a change in stimulation parameters were preceded by a 5-second linear ramp-down at carrier frequency and followed by a 5-second linear ramp-up at carrier frequency. Dedicated ramping windows were included in the data acquisition protocol to mark these periods for subsequent exclusion from analysis. Transitions between blocks of the same condition type did not involve ramping.

##### Electrode montage

Circular rubber electrodes (1.5 cm diameter) were positioned at 10-10 positions F7, F8 (frontal pair) and P5–P6 (parietal pair). Electrode sites were prepared using NuPrep exfoliating gel (Weaver and Company, Aurora, CO, USA) to abrade the skin surface, followed by alcohol wipes and gauze to remove debris. Electrodes were affixed using Ten20 conductive paste (Weaver and Company, Aurora, CO, USA) and stabilised with microporous surgical tape. Impedance at each electrode was verified to be below 5 kΩ using an impedance checker (EL-CHECK, BIOPAC Systems, Inc., Goleta, CA, USA).

##### Stimulation perception thresholds

We determined perceptual thresholds using conventional transcranial alternating current stimulation (tACS) and TIS.

Frist 5Hz tACS was delivered separately to each electrode pair, with increasing intensity until the participant reported a sensation. Second, temporal interference stimulation (df=5Hz) was delivered with increasing amplitude until sensation was reported or the 3mA maximum was reached. At each step, participants were asked to report the presence and quality of any sensation (tingling, itching, warmth, or pressure) and individual thresholds were recorded. All participants tolerated 2mA temporal interference stimulation without reporting any sensation.

##### Stimulation waveform recording and timing

TIS protocols were pre-programmed using the TIBS-R Scripter software. During the nap, a custom MATLAB script-controlled stimulus timing by sending TTL triggers through a National Instruments USB-6343 data acquisition (DAQ) device to the TIBS-R device. Identical stimulation parameters were simultaneously routed from a second pair of stimulator channels to the DAQ device, where the stimulation waveform was recorded at 20 kHz via a custom C++ function using Lab Streaming Layer (LSL). Event markers identifying the condition, block number, and trial number were transmitted as LSL string markers from the MATLAB control script.

##### Side effects monitoring

We included a post-nap questionnaire administered immediately after waking to monitor side-effects. Participants rated potential stimulation-related sensations — itching, tingling, headache, scalp pain, neck pain, burning, warmth or heat, metallic taste, nervousness or anxiety, discomfort, dizziness, nausea, and other — on a six-point scale (0=no sensation; 1=very weak; 2=weak; 3=moderate; 4=strong; 5=severe). For any reported sensation, participants additionally indicated the timing of onset (beginning, middle, or end of the nap; throughout the nap; or after waking), and the anatomical localisation (diffuse; localised to the right or left side; close to the left or right ear; close to the forehead; or close to the back of the head).

##### EEG, EOG, EMG recording

Scalp EEG was recorded using 30 passive Ag/AgCl electrodes mounted in an EasyCap EEG cap (EasyCap GmbH, DE) at 10–20 positions: Fp1, Fp2, F7, F3, Fz, F4, F8, FC5, FC1, FC2, FC6, T7, C3, Cz, C4, T8, TP9, CP5, CP1, CP2, CP6, TP10, P7, P3, Pz, P4, P8, POz, O1, and O2. Linked mastoids (TP9 and TP10) served as reference and AFz as ground.

Electrooculography (EOG) was recorded as bipolar signal, using two electrodes positioned 1 cm lateral and 1 cm superior to the left and right outer canthi (LOC and ROC), each referenced to the contralateral mastoid (LOC to right mastoid, ROC to left mastoid). Submental electromyography (EMG) was recorded as bipolar signal from a single chin electrode referenced to a mastoid electrode.

All EXG electrodes had an impedance below 20 kΩ, the ground electrode below 5 kΩ.

All physiological data was recorded using an actiCHamp Plus amplifier (Brain Products GmbH, Gilching, Germany) at 10 kHz sampling rate.

EEG data, stimulation waveforms and stimulation event markers were recorded via LSL, and raw data were stored in XDF (Extensible Data Format) files.

### EEG analysis

#### Outcome measures

The primary outcome measure was spindle density during N2 sleep, computed in the 2-second stimulation epochs. We hypothesised that 5Hz temporal interference stimulation would selectively increase spindle density relative to both the 1Hz and OFF conditions. Secondary outcome measures included spindle-band spectral power (12–16Hz), spindle amplitude, spindle duration, and spindle frequency. Additionally, we examined the temporal dynamics of spindle occurrence across the 2-second stimulation window and the probability of spindle occurrence per trial using generalised linear mixed-effects models.

#### EEG Preprocessing

##### Data import

All preprocessing was performed in MATLAB (R2023a, The MathWorks, Natick, MA) using the FieldTrip toolbox (Oostenveld et al., 2011) and a custom-built codebase.

Initially, three data streams were extracted from each XDF file: the EXG stream (10 kHz), event markers (LSL string markers), and the TIS waveform (20 kHz). XDF files were loaded using the *load_xdf* function and converted to FieldTrip-compatible data structures using *xdf2fieldtrip*. Timestamps across all three streams were aligned to the start of the recording. Auxiliary channels (EMG, left EOG, right EOG) were separated from the main EEG channels and processed independently before being re-merged after the main EEG pipeline. Electrode labels and positions were added from an EasyCap provided layout file (BrainCap) with 32 scalp positions.

##### Artefact detection and interpolation

A custom slope-based artefact detection algorithm targeted high-amplitude transient artefacts occurring during stimulation ramping. Data were first downsampled to 500Hz with automatic anti-aliasing, and bandstop-filtered (4th-order Butterworth, zero-phase, 47.5– 52.5Hz). The channel with the highest variance of its absolute first-order temporal derivative was selected as the artefact detection channel.

Artefact detection was restricted to temporal windows surrounding ramping events (ramp onset − 1s to next marker + 1 s). The absolute first-order derivative was smoothed with an 11-sample moving median filter, and the detection threshold was set at the median plus 15 times the median absolute deviation (MAD).

Detected artefact samples were dilated by 200ms on each side. Artefact segments up to 2000ms in duration were interpolated by a linear fit between flanking samples, applied independently per channel. On average, 0.53% of data (SD=0.34%) were rejected across the total recording time.

##### Filtering

After artefact interpolation, a three-stage zero-phase filtering cascade was applied to the continuous data using MATLAB’s *filtfilt* (forward-backward filtering). First, a high-pass filter (2nd-order Butterworth, effective 4th-order via *filtfilt*, 0.3Hz cutoff) removed slow drifts. Second, a low-pass filter (4th-order Butterworth, effective 8th-order, 30Hz cutoff) attenuated high-frequency noise. Third, a notch filter (IIR, Q=10, 50Hz) removed residual line noise. Filters were applied sequentially per channel in the order: high-pass, low-pass, notch.

##### Resampling

After filtering, both main EEG and auxiliary channel data were resampled to 500Hz using FieldTrip’s *ft_resampledata* with detrending disabled.

##### Automatic channel rejection

Noisy channels were identified automatically using a variance-based criterion computed over the stimulation interval (from *STIM_START* to *STIM_STOP* markers). Per-channel variance was log-transformed and z-scored across channels. Channels with z-scored log-variance exceeding 1.5 were flagged for rejection (mean=3.50 ± 0.83 channels), with the majority of rejections being EEG electrodes on close to stimulation electrodes (F7 and F8 in 8/20 participants and 12/20 participants, P7 and P8 in 11/20 and 14/20 participants respectively).

##### Channel interpolation

Rejected channels were reconstructed using FieldTrip’s nearest-neighbour weighted interpolation (*ft_channelrepair*, method=‘nearest’). Electrode positions were loaded from the EasyCap layout file. Channel neighbourhoods were defined using distance-based adjacency (*ft_prepare_neighbours*, method=‘distance’), with the neighbour distance threshold set adaptively to twice the median nearest-neighbour distance in the electrode configuration.

##### Re-referencing

All EEG channels were re-referenced to the average of the linked mastoids (TP9 and TP10) using FieldTrip’s *ft_preprocessing*. Reference channels were retained in the data.

#### Segmentation and trial rejection

We restricted our analysis to N2 sleep, which matched our model parametrisation and where the highest sigma power and sleep spindle event density occurs during NREM sleep^1^.

Continuous data were segmented into trials based on event markers. Each trial was assigned to one of three condition labels: OFF, 1Hz, 5Hz (stimulation epochs), plus ramping windows. Trials were further subdivided into PRE [(−1) - 0 s], STIM [0 - 2 s] and POST [2 - 5 s] windows for some analyses.

Epochs were classified as artefact by 1) automatically checking for temporal overlap between each epoch’s boundaries and the artefact segments identified during the slope-based detection step, 2) and manual rejection of the first trial of each OFF block (following a ramping-down transition). These first OFF trials contained artefacts from the combined effects of the ramping-down and the stimulator’s transition from 2000Hz (0 mA) to 0Hz (0 mA). This transition produced a brief artefact that was not fully attenuated by the hardware filters. This exclusion resulted in slightly fewer trials in the OFF condition relative to the active stimulation conditions.

From an initial 285.6 ± 107.7 stimulation epochs per participant (total: 5,711), 11.4 ± 3.2% were excluded: 7.2% due to artefact overlap with slope-based detection segments and 4.0% due to first-OFF-trial exclusion. Restriction to N2 sleep excluded a further 47.7% of surviving epochs. After all exclusions, the mean number of clean N2 trials per participant was 47.7 ± 25.3 for 5Hz (954 total), 45.8 ± 27.5 for 1Hz (915 total), and 39.2 ± 19.5 for OFF (784 total), yielding 2,653 stimulation epochs across all 20 participants.

#### Sleep Scoring

A continuous version of the recorded of frontal (F3, F4), central (C3, C4), parietal (P3, P4), and occipital (O1, O2) derivations, plus EOG and EMG channels were exported as European Data Format (EDF) file resampled to 256Hz. Sleep staging was performed manually according to American Academy of Sleep Medicine (AASM, 2017) criteria using Domino software (Somnomedics GmbH, Randersacker, DE). The data was segmented into 30-second epochs and classified as Wake, N1, N2, N3, or REM based on standard criteria by a trained sleep researcher blind to stimulation conditions.

### Spectral Analysis

#### Additional optional Trial selection

Before spectral analysis trials underwent a multi-stage quality control pipeline. First, per-channel trial power was assessed using a robust z-score (|x − median| / [1.4826 × MAD]), and trials exceeding a MAD threshold of 3.0 were rejected. Second, a z-scored spindle power inclusion filter was applied to retain only trials exhibiting meaningful spindle-band activity. For each trial, mean power in the sigma band (12-16Hz) was averaged across a −1 to 3s time window (relative to stimulation onset) separately for each electrode. Power values were then z-scored across trials within each electrode and participant. Trials were retained if the z-score exceeded 2.0 in at least one channel. Third, power values were converted to decibels (10 × log_10_). Trial indices surviving the primary topographic analysis were saved and reused across all power figure panels to ensure consistency.

#### Sigma power

Sigma power was computed for each trial using the multitaper spectral analysis (using Fieldtrip’s ft_freqanalysis with mtmconvol). For electrode-level analyses, a multitaper spectrum was computed for each trial in a 0.50–1.50 second time window following stimulation onset. The mean log-transformed raw sigma power (12–16Hz) was then averaged across trials within each condition.

#### Time-frequency representation

Time-frequency representations (TFRs) were computed using multitaper spectral analysis. A TFR was computed for each trial (using FieldTrip *ft_freqanalysis* with *mtmconvol*). TFRs were computed with the following parameters: frequency range 0.5–30Hz in 0.25Hz steps, a fixed time window of 1.0 second, spectral smoothing of 1.0Hz via discrete prolate spheroidal sequence (DPSS) tapers, time step of 0.05 seconds (50 ms), pre- and post-trial padding of 6.0 seconds each, and zero-padding to the next power of 2.

#### Sigma power topography

Topographic maps of sigma power were computed by averaging TFR power across the 12– 16Hz frequency window and the 0.50–1.50 second post-onset time window for each electrode, condition, and participant. Condition differences were assessed using cluster-based permutation testing (see *Statistical Analysis*).

#### Spindle event analysis

Sleep spindles were detected algorithmically using the YASA toolbox (Vallat & Walker, 2021) implemented in Python, applied to the 256Hz EDF exports. Detection was performed on all 30 EEG channels using *yasa.spindles_detect* with the following parameters: spindle frequency band of 10–16Hz and duration range of 0.3–3.0 seconds. All other detection parameters were left at YASA’s internal defaults: broadband filter of 1–30Hz, root-mean-square (RMS) moving window of 0.3 seconds, relative sigma power threshold of 0.2, Pearson correlation threshold between the raw and sigma-filtered signal of 0.65, RMS amplitude threshold of the mean plus 1.5 standard deviations of the sigma RMS envelope, and minimum inter-spindle distance of 0.5 seconds for event merging.

The YASA detection algorithm operates as follows: (1) the EEG signal was bandpass-filtered in the sigma band (10–16Hz) and broadband (1–30Hz) using FIR filters; (2) the RMS of the sigma-filtered signal was computed using a 0.3-second moving window; (3) relative sigma power was calculated as the ratio of sigma RMS to broadband RMS; (4) candidate events were identified where relative sigma power exceeded 0.2, correlation between the raw and sigma-filtered signal exceeded 0.65, and sigma RMS exceeded 1.5 standard deviations above the mean; (5) events separated by less than 0.5 seconds were merged; (6) events were retained only if their duration fell within 0.3–3.0 seconds; and (7) peak frequency was estimated via FFT for each detected event. For each spindle event YASA computed onset time, peak time, end time, duration, amplitude, RMS, absolute power, relative power, peak frequency, number of oscillations, symmetry index, and channel label.

For all spindle event analyses, we only included spindle events satisfying the following criteria: (1) peak frequency between 12 and 16Hz, (2) peak-to-peak amplitude between 15 and 100 µV, (3) duration between 0.5 and 3.0 seconds.

The final set of spindle events was assigned a sleep stage based on spindle onset relative to 30-second sleep stage annotations and assigned to experimental conditions against trial boundaries derived TTL markers. Subject-level means were computed per condition by averaging across all detected spindles (meeting inclusion criteria) across all electrodes during N2 sleep.

Grand-average spindle waveforms were computed per condition for visualisation. EEG data were extracted in a ±0.5 second window around each detected spindle peak at a central ROI (including electrodes: FC1, FC2, C3, Cz, C4, CP1, CP2). Per-event waveforms were interpolated to a common 200-point time axis, averaged within participant (minimum 20 events required), and then averaged across participants. Waveforms were displayed as mean ± standard error of the mean (SEM).

##### Spindle density

Spindle density was computed for each participant, condition, and electrode, as the number of spindles with onset during STIM windows ([0-2 s]) divided by the total duration of STIM windows in minutes. Density was computed at the electrode level; for analyses requiring a single density value per participant per condition, electrode-level densities were averaged across all 30 electrodes. For mixed-effects models retaining electrode-level data, electrode was modelled as a nested random effect within participant.

##### Spindle density topography

Group-level topographic maps of spindle density were generated by averaging electrode-level densities across participants for each condition. Differences between condition were assessed using cluster-based permutation statistics (see *Statistical Analysis*).

##### Trial-based histogram of spindle onsets

The temporal distribution of spindle onsets relative to trial onset was analysed using binned histograms. The time window was −0.5 to 2.0 seconds relative to trial onset, divided into five 0.5-second bins (bin centres: −0.25, 0.25, 0.75, 1.25, 1.75 seconds). For each electrode, the spindle rate in each bin was computed as the number of spindle onsets falling within the bin divided by the number of N2 trials for that participant–electrode–condition combination. Subject-level values were obtained by averaging across electrodes.

### Statistical Analysis

All statistical analyses were performed in MATLAB (R2023a) and RStudio with R (v4.5.2). Significance was assessed at α=0.05 (two-tailed) unless stated otherwise.

#### Cluster-based permutation testing

Topographic condition differences in spindle-band power and spindle density were assessed using cluster-based permutation testing. The procedure was as follows: (1) Per-electrode paired t-tests were computed between conditions (within-subject differences); (2) Electrodes exceeding the cluster-forming threshold (p < 0.15) were identified. A liberal cluster-forming threshold was chosen because sleep spindle modulation effects were expected to be spatially distributed across multiple electrodes rather than sharply localised, and a more stringent threshold (e.g., p < 0.05) risks fragmenting a single diffuse effect into multiple small clusters or failing to detect it altogether^34^. Family-wise error control is maintained at the cluster level through the permutation procedure, rendering the cluster-forming threshold a sensitivity parameter rather than an inferential threshold. (3)Spatially adjacent suprathreshold electrodes were grouped into clusters using distance-based adjacency (FieldTrip *ft_prepare_neighbours*, distance method, threshold=0.25 in normalised units). (4)The cluster statistic was computed as the sum of absolute t-values within each cluster. Clusters smaller than 2 electrodes were discarded. (5)A null distribution of maximum cluster statistics was generated by 5000 permutations of subject-level random sign-flips of within-subject difference maps.

(6) Cluster p-values were computed as the proportion of permutation maximum cluster statistics exceeding the observed cluster statistic (one-tailed). Clusters were considered significant at p < 0.05.

For time-frequency analyses, FieldTrip’s built-in cluster-based permutation framework (*ft_freqstatistics*) was used with dependent-samples t-tests, cluster-forming α=0.15, maximum-sum cluster statistic, two-tailed testing, and 5000 permutations.

#### Linear mixed-effects models

Linear mixed-effects models (LMMs) were used for continuous outcome variables (spindle density, spindle-band power, spindle duration, amplitude, and frequency). Models were fitted using restricted maximum likelihood (REML) estimation via MATLAB’s *fitlme*.

For spindle density, the primary model was specified as:

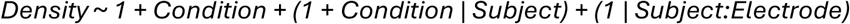

where Condition (5Hz, 1Hz, OFF) was the fixed effect of interest, with random intercepts and slopes for Condition within Subject and random intercepts for Electrode nested within Subject. This maximal random-effects structure converged in all use cases. Model fit was evaluated using marginal and conditional R^2^ (Nakagawa & Schielzeth approximation).

For spindle metrics (duration, amplitude, frequency), the same model formula was used:

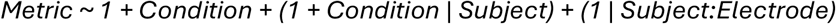

Omnibus F-tests were computed via analysis of variance on the LMM.

#### Generalised linear mixed effects models

In addition to density analyses, trial-based spindle probability (i.e. a spindle was present vs. absent within each 2-second Condition window) was analysed using generalised linear mixed-effects models (GLMMs) with a binomial distribution and logit link function.

The condition window was split into four 0.5s time bins (plus a pre window of −0.5 to 0s prior to stimulation start) to distinguish spindle timing within trials. The binary outcome was spindle occurrence (0/1) per observation (participant x electrode x trial x time bin).

The GLMM analyses were conducted in R using the *lme4* package^35^. Model diagnostics were assessed using the DHARMa package (simulated residuals with 500 simulations, uniformity tests, dispersion tests, zero-inflation tests, and bootstrap outlier detection).

We compared six models of increasing complexity. (M0) Condition as the sole fixed effect with random intercepts for Subject and Electrode; (M1) main effects of Condition and Time Bin with a random intercept for Subject; (M2) adding the Condition x Time Bin interaction; (M3) adding a random intercept for Electrode; (M4) adding a random intercept for the Subject x Electrode combination; and (M5) including random slopes for Condition by Subject together with a random intercept for Electrode. All models were fitted via maximum likelihood (Laplace approximation) using the BOBYQA optimizer. Model selection was based on the Akaike Information Criterion (AIC), Bayesian Information Criterion (BIC), and likelihood ratio tests for nested models, as implemented in the performance package^36^. Significance in all models was assessed with Type III Wald chi-square tests.

#### Post-hoc pairwise comparisons

Planned pairwise comparisons between conditions (5Hz vs. OFF, 1Hz vs. OFF, 5Hz vs. 1Hz) were computed using linear contrasts on fixed effect coefficients with standard errors derived from the coefficient covariance matrix. For LMMs, contrasts were evaluated using F-tests. For GLMMs, Wald z-tests were used, and results were expressed as odds ratios with 95% Wald confidence intervals on the log-odds scale. Multiple comparison corrections were applied using the Holm-Bonferroni step-down procedure across the three pairwise comparisons per analysis. For analyses involving multiple time bins, false discovery rate (FDR) correction (Benjamini-Hochberg, positive dependency assumption) was applied within each contrast pair across bins.

#### Effect sizes

Effect sizes were reported as Cohen’s d for continuous outcomes. For LMMs, Cohen’s d was computed as the contrast estimate divided by the pooled standard deviation (square root of the sum of subject and residual variance components). For GLMMs, Cohen’s d was converted from the log odds ratio using the formula: d=log(OR) × √3 / π ^37^. Odds ratios with 95% confidence intervals were additionally reported for all GLMM contrasts. For temporal histogram analyses, Cohen’s h (arcsine transformation for proportion differences) was used as an effect size measure.

### Software

EEG preprocessing and analysis were performed in MATLAB (R2023a) using the Signal Processing Toolbox, and the Statistics and Machine Learning Toolbox, FieldTrip^38^ for spectral analysis, topographic plotting, and cluster-based permutation statistics; and EEGLAB^39^. Automated spindle detection was performed in Python (v3.7) using MNE-Python (v. 1.20) and YASA (v.0.7.0^20^).

Sensitivity analyses were conducted in R using *lme4* for GLMM fitting, *car* for Type III tests, *emmeans* for estimated marginal means and contrasts, *DHARMa* for model diagnostics, and *performance* for model comparison.

## Data and code availability

All custom code central to the conclusions in this paper is publicly available.

Simulation and analysis scripts related to the computational model are in the public GitHub repository https://github.com/bazhlab-ucsd/ti-thalamic-stim-public.git.

Data analysis code: https://github.com/BartschLab/Spindle-TI

Experimental data: The EEG dataset has been deposited on OpenNeuro in BIDS format. For peer review, access is provided via an anonymous OpenNeuro reviewer link submitted separately through the journal submission system. The public OpenNeuro DOI will be added upon acceptance/public release.

## Grant/Support

MGNZ, MB: 1R01AG099626, 1R01MH125557, 1RF1NS132913, TR, PO, UB, IV: BB/Y011856/1

UB, DJD: UK Dementia Research Institute [award number UKDRI-7206] through UK DRI Ltd, principally funded by the UK Medical Research Council, and additional funding partner Alzheimer’s Society.

DJD: NIHR Oxford Health Biomedical Research Centre [NIHR 203316].

## Author contributions

### CRediT

MGNZ: Methodology, Software, Formal analysis, Visualization, Investigation, Writing - Original Draft, Writing - Review & Editing; TR: Methodology, Software, Validation, Data Curation, Formal analysis, Visualization, Investigation, Writing - Original Draft, Writing - Review & Editing; PO: Investigation, Formal analysis; EN: Resources, NK: Resources; DJD: Writing - Review & Editing, Funding acquisition; IV: Writing - Review & Editing, Resources, Supervision, Project administration, Funding acquisition; MB: Conceptualization, Supervision, Project administration, Funding acquisition, Writing - Review & Editing; UB: Conceptualization, Supervision, Project administration, Funding acquisition, Writing - Original Draft, Writing - Review & Editing,

