## Supplementary Information for "Resonance-driven enhancement of sleep spindles using thalamic temporal interference stimulation"

### Figure S1

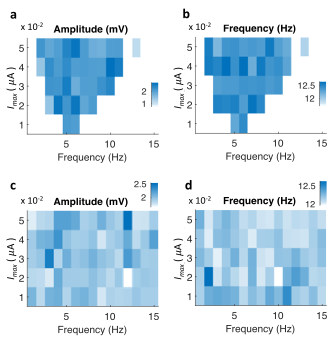

**Figure S1. Spindle event properties in the model in response to stimulation**

Spindle amplitude (a, c) and frequency (b, d) in the stimulation frequency-intensity sweep for models without cortical inputs to the thalamus (a, b) or with intact full connectivity (c, d). Amplitude and frequency do not systematically vary with stimulation parameters.

### Figure S2

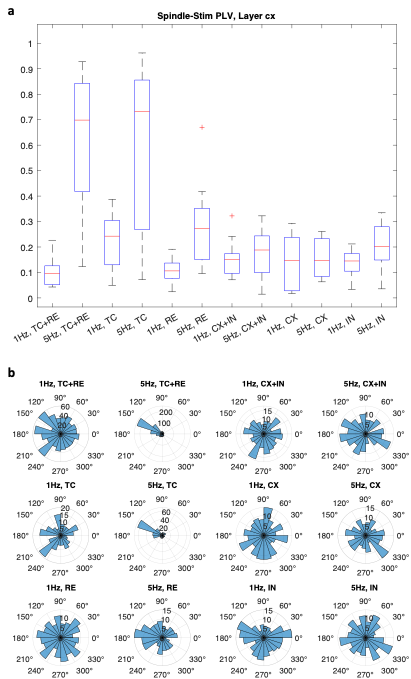

**Figure S2. Stimulating other cell populations in the model.**

(a) Phase locking value (PLV) between spindle onsets and periodic identical stimulation delivered to different cell populations. Simulations were 180s with 2s 1Hz or 5Hz sinusoidal input applied every 8 seconds (6s refractory period). Target populations are indicated in the x axis, with highest intensity at the network center. High PLV indicates high entrainment of spindle onsets to stimulation timing. (b) Histograms of spindle event phase with respect to stimulation period for each target population and stimulation frequency. Alignment between spindles and stimulation onsets is strongest for TC or TC+RE target populations.

### Figure S3 CONSORT diagram

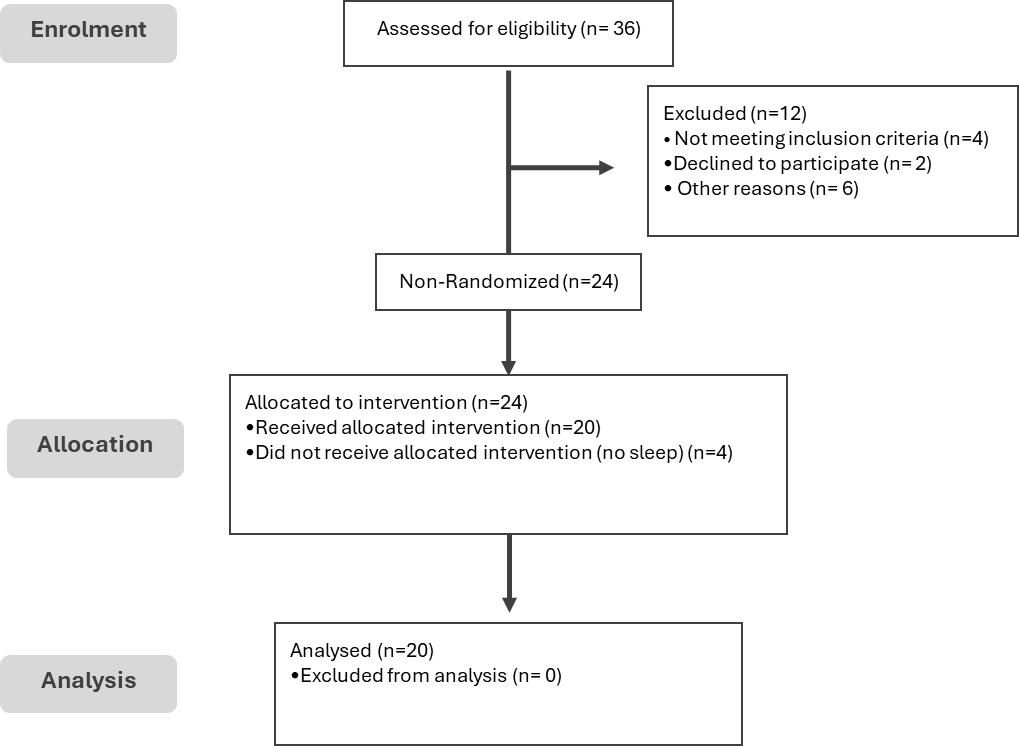

***Figure S3 | Consort diagram****: Recruitment for the TIS-EEG study.*

### Figure S4

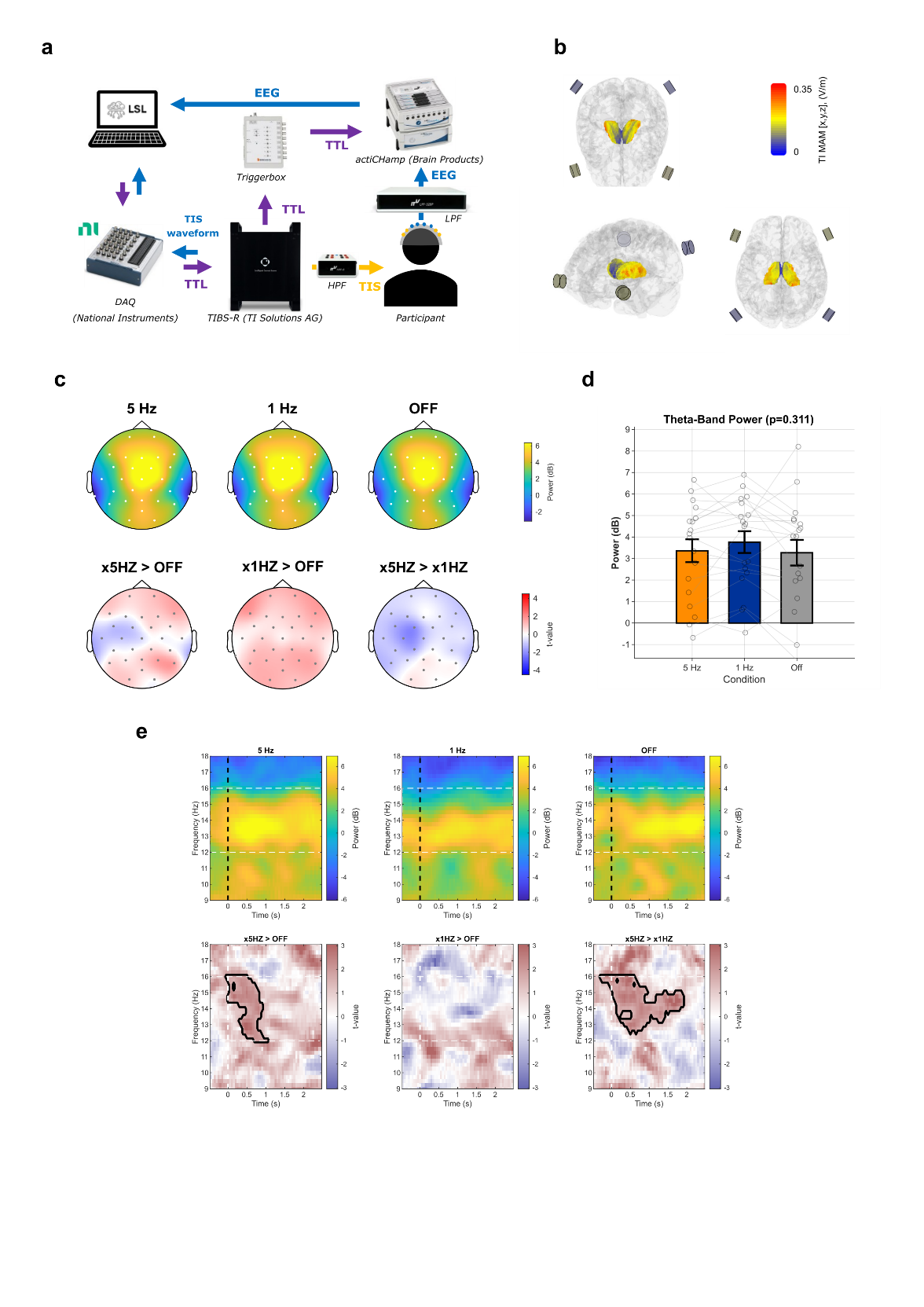

***Figure S4 | Experimental setup, electric field modelling and control analyses.***
*(a) Schematic of the data acquisition setup. EEG was recorded from the participant via a hardware low-pass filter (LPF) and an actiCHamp amplifier (Brain Products). Temporal interference stimulation (TIS) was delivered to the participant via a TIBS-R stimulator (TI Solutions AG), with triggers routed through a triggerbox to the amplifier. The TIS waveform was simultaneously recorded via a data acquisition device (DAQ; National Instruments). All data streams were synchronised and recorded on a PC via Lab Streaming Layer (LSL).*

*(b) Predicted EM field intensity in the thalamus for one example participants. Simulated electric field distribution of the TIS montage, modelled using Sim4Life. Colour scale indicates the maximum amplitude modulation (TI MAM) of the electric field (V/m), electrode positions are overlaid.*

*(c) Upper row: scalp topographies of theta-band power (4–8 Hz, dB) during stimulation for each condition. Lower row: pairwise condition differences displayed as t-value maps. No significant clusters were observed (cluster-based permutation test; cluster-forming threshold p < 0.15, cluster-level α = 0.05). d, Global average theta-band power (4–8 Hz) across conditions. Bar plot error bars indicate ± 1 SEM. No significant effect of condition was observed (linear mixed-effects model; p = 0.311).*

*(e) Upper row: time–frequency spectrograms at electrode CP5 for each condition during the stimulation window. Dashed white horizontal lines indicate the spindle frequency band (12–16 Hz); dashed black vertical lines indicate stimulation onset. Lower row: pairwise condition differences displayed as t-value maps. Black contours outline significant clusters (cluster-based permutation test), revealing greater power in the 5 Hz condition compared with both OFF and 1 Hz.*

### Figure S5

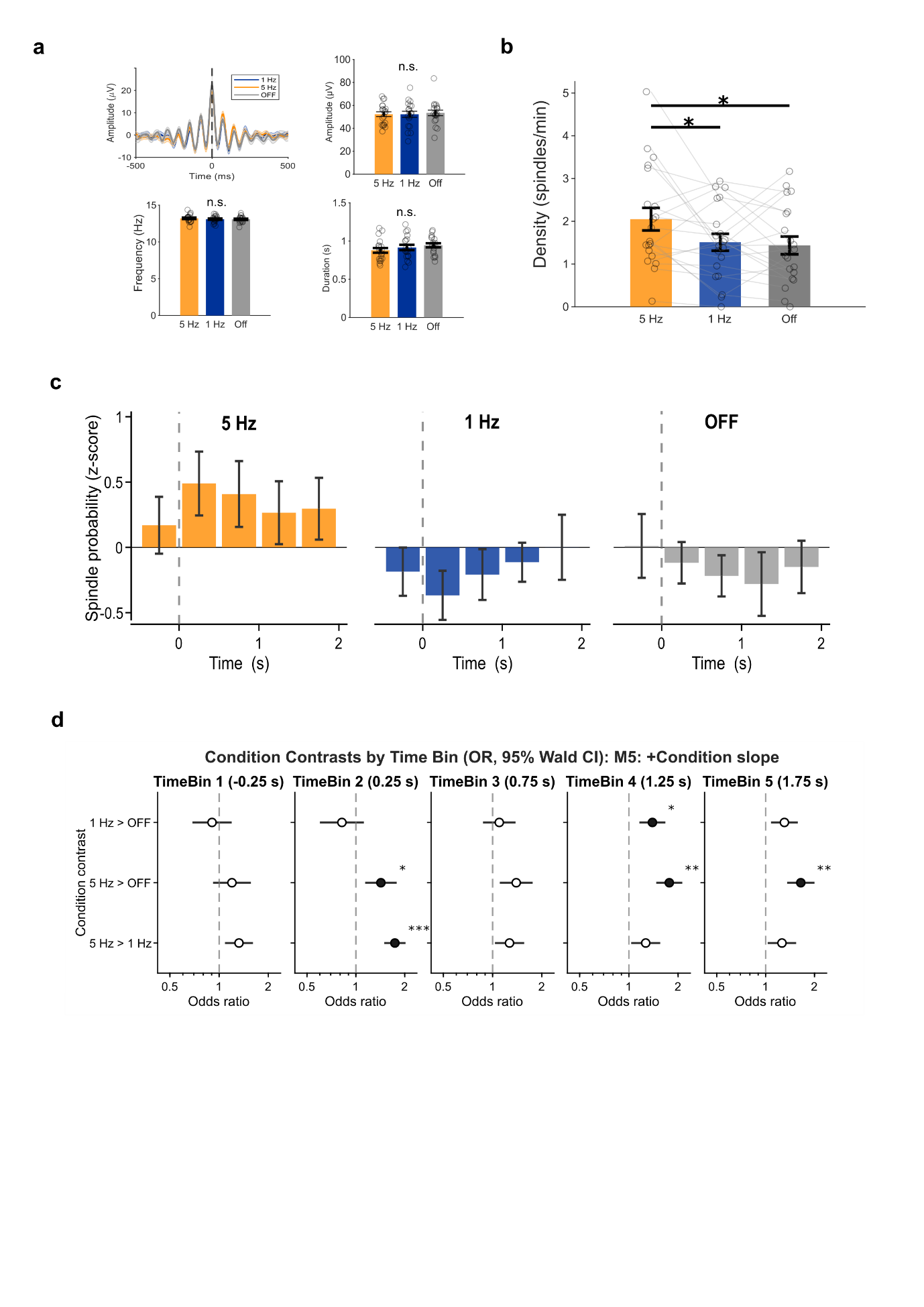

***Figure S5 | Time-resolved spindle probability and pairwise condition contrasts.***
*(a) Detected spindle events did not differ in waveform (top left) or in derived properties such as amplitude (tope right), frequency (bottom left) or duration (bottom right) across conditions (Table S12, S13)*

*(b) Spindle density (events/min) was higher in the 5Hz condition compared to 1Hz and OFF condition (Linear mixed model analysis, Table S14, S15))*

*(c) Spindle probability per condition resolved into 0.5 s time bins relative to stimulation onset (dashed vertical line), expressed as z-scored probability (normalised across all bins and conditions). Error bars indicate ± 1 SD.*

*(d) Odds ratios (OR) with 95% Wald confidence intervals for pairwise condition contrasts at each time bin, derived from the generalised linear mixed model with time bin and condition (M5). Each row represents a condition contrast (5 Hz > 1 Hz, 5 Hz > OFF, 1 Hz > OFF). The dashed vertical line at OR = 1 indicates no difference. Filled circles denote significant contrasts; open circles denote non-significant contrasts. Significant effects include 5 Hz > 1 Hz and 5 Hz > OFF at the 0–0.5 s bin, 5 Hz > OFF and 1 Hz > OFF at the 1–1.5 s bin, and 5 Hz > OFF at the 1.5–2 s bin. All panels: n = 20 participants. *p < 0.05, **p < 0.01****, *******p < 0.001, Table S21)*

### Figure S6

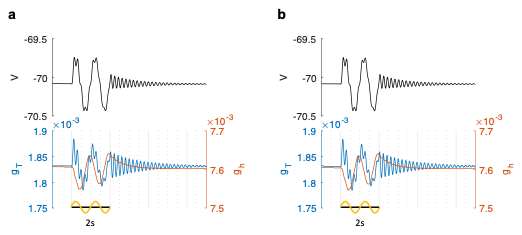

***Figure S6 | 1Hz stimulation in minimal computational models.***

*(a) 1Hz, 2s,* 0.01 $\mu$A *sub-threshold stimulation (sine wave) for an isolated TC cell in the bistable regime (GT=2.34, Gh=0.016). From top to bottom: membrane voltage, calcium influx, T current and h current. No spikes are elicited. (d) as (c), for one TC cell in a minimal circuit with 2 RE and 2 TC interconnected cells. Stimulation amplitude is identical to Figure 3c, d.*

**Participant demographics**

|  | **Recruited (N = 24)** | **Final sample (N = 20)** |
| --- | --- | --- |
| **Age: Mean (SD) , [range]** | 27.5 (SD = 4.95) , [19-35] | 27.0 (SD = 4.94) , [19-35] |
| **Sex: (M/F)** | 15 / 9 | 12 / 8 |

### Sleep architecture analysis

Table S 1: Stimulation conditions per sleep stage

| Stage | 5Hz | 1Hz | OFF |
| --- | --- | --- | --- |
| N1 | 11.4 ± 9.6 | 12.5 ± 10.3 | 10.1 ± 10.6 |
| N2 | 60.3 ± 25.6 | 57.9 ± 28.2 | 60.8 ± 26.9 |
| N3 | 24.0 ± 25.5 | 24.3 ± 25.8 | 24.5 ± 25.8 |
| REM | 4.3 ± 9.0 | 5.3 ± 11.7 | 4.6 ± 7.3 |

Table S 2: Repeated measures ANOVA per stage across conditions:

| Stage | F | df | p |
| --- | --- | --- | --- |
| N1 | 0.64 | 2, 38 | 0.55 |
| N2 | 0.83 | 2, 38 | 0.44 |
| N3 | 0.18 | 2, 38 | 0.83 |
| REM | 0.41 | 2, 38 | 0.67 |

Table S 3: Wake epochs per stimulation condition

(Number of wake epochs in between the first N2 epoch and the last sleep epoch, per condition)

| Condition | Mean | SD |
| --- | --- | --- |
| 5Hz | 7.25 | 11.92 |
| 1Hz | 7.40 | 11.86 |
| OFF | 5.70 | 10.26 |

Table S 4: Repeated measures ANOVA per stage across conditions:

| Source | SS | Df | MS | F | P | P (GG) |
| --- | --- | --- | --- | --- | --- | --- |
| Condition | 35.43 | 2 | 17.72 | 1.65 | .21 | .21 |
| Error (residual) | 409.23 | 38 | 10.77 |  |  |  |
| Total | 444.67 | 40 |  |  |  |  |

### Spectral Analysis

#### Power spectrum

Table S 5: Cluster corrected, univariate test of absolute power

One-sample t-statistic on within-subject differences (equivalent to paired t-test).

For each condition pair, within-subject differences are computed per frequency, one-sample t-test performed for each frequency. Frequencies with p<0.15 are grouped into frequency adjacent clusters, and cluster masses compared through 5000 permutations.

With subject-level sign-flip permutations (N=5000): cluster forming threshold 0.15, 1-20Hz

Final cluster alpha: <.05

| **Contrast** | **Cluster mass value** | **Cluster f range** | **p-value** | **Cohen’s d** |
| --- | --- | --- | --- | --- |
| **5Hz > 1Hz:** |  |  |  |  |
| Null distribution 95^th^ percentile | 15.66 |  |  |  |
| **Cluster 1** | **29.42** | **10.74-16.60Hz** | **0.010** | **0.58** |
| **5Hz > OFF**: |  |  |  |  |
| Null distribution 95th percentile | 16.37 |  |  |  |
| **Cluster 1** | **26.16** | **10.74-16.60Hz** | **0.017** | **0.53** |
| **1Hz = OFF** |  |  |  |  |
|  |  |  | ns |  |

Table S 6: Cluster descriptives

Mean +/- SD (dB) of identified clusters

| **Condition** | **Mean** | **SD** |
| --- | --- | --- |
| **5Hz** | 1.92 | 2.53 |
| **1Hz** | 1.30 | 2.73 |
| **OFF** | 1.26 | 2.23 |

#### Fast spindle power topography

Table S 7 Average spindle-band power across all electrodes

| **Condition** | **Mean (dB)** | **SD (dB)** |
| --- | --- | --- |
| 5Hz | 0.01 | 2.79 |
| 1Hz | -1.00 | 0.60 |
| OFF | -1.27 | 0.57 |

Table S 8 Cluster corrected, univariate test results

> One-sample t-statistic on within-subject differences (equivalent to paired t-test).

For each condition pair, within-subject differences are computed per electrode, one-sample t-test performed for each electrode. Electrodes with p<.15 are grouped into spatial clusters, and cluster masses compared through 5000 permutations.

Subject-level sign-flip permutations (n=5000), cluster threshold 0.15, minimum size = 2 electrodes

| **Contrast** | **Cluster mass value** | **Cluster included electrodes** | **p-value** | **Cohen’s d** |
| --- | --- | --- | --- | --- |
| **5Hz > 1Hz:** |  |  |  |  |
| Null distribution 95^th^ percentile | 37.31 |  |  |  |
| **Cluster 1** | **64.83** | **26/30** | **0.009** | **0.70** |
| **5Hz > OFF**: |  |  |  |  |
| Null distribution 95th percentile | 39.87 |  |  |  |
| **Cluster 1** | **55.85** | **23/30** | **.018** | **0.73** |
| **1Hz > OFF** |  |  |  |  |
|  |  |  | ns | 0.11 (global) |

#### Theta power topography stats

subject-level sign-flip permutations (5000) , cluster threshold 0.15, minimum size = 2 electrodes

Table S 9 Average theta band (4-8Hz) power across all electrodes (dB):

| Condition | Mean | SD |
| --- | --- | --- |
| 5Hz | 3.58 | 2.38 |
| 1Hz | 3.83 | 2.11 |
| OFF | 3.27 | 2.53 |

Table S 10 Cluster corrected, univariate test results

| **Contrast** | **Cluster mass value** | **Cluster included electrodes** | **p-value** | **Cohen’s d** |
| --- | --- | --- | --- | --- |
| **5Hz > 1Hz:** |  |  |  |  |
| Null distribution 95^th^ percentile | 34.31 |  |  |  |
| Cluster 1 | **3.90** | **2/30** | p = 0.439 | **-0.19)** |
| **5Hz > OFF**: |  |  |  |  |
| Null distribution 95th percentile | 31.09 |  |  |  |
| Cluster 1 | **3.29** | **2/30** | p = 0.568 | **0.08** |
| **1Hz = OFF** |  |  |  |  |
|  |  |  | ns | 0.30 |

#### Stimulation triggered multi-taper spectrogram at electrode CP5 (time-frequency analysis)

Table S 11 Cluster corrected, univariate test results

| **Contrast** | **Cluster mass value** | **Cluster time range** | **Cluster f range** | **p-value** | **Cohen’s d** |
| --- | --- | --- | --- | --- | --- |
| **5Hz > 1Hz:** |  |  |  |  |  |
| Null distribution 95^th^ percentile | 307.82 |  |  |  |  |
| **Cluster 1** | 317.02 | -0.05 – 1.90s | 13.2-16.0Hz | 0.047 | 0.54 |
| **5Hz > OFF**: |  |  |  |  |  |
| Null distribution 95th percentile | 275.07 |  |  |  |  |
| **Cluster 1** | 323.36 | -0.25 - 1.00s | 12.0-15.3Hz | 0.031 | 0.69 |
| **1Hz = OFF** |  |  |  |  |  |
|  |  |  |  | ns |  |

### Spindle event analyses

#### Spindle event properties

Table S 12: Spindle event properties

|  | **Spindle duration (s)** | |
| --- | --- | --- |
| **Condition** | **Mean** | **SD** |
| 5Hz | 0.87 | 0.13 |
| 1Hz | 0.91 | 0.15 |
| OFF | 0.94 | 0.11 |
|  | **Spindle amplitude (µV):** | |
|  | **Mean** | **SD** |
| 5Hz | 51.31 | 8.85 |
| 1Hz | 52.43 | 11.70 |
| OFF | 53.56 | 11.00 |
|  | **Spindle frequency (Hz)** | |
|  | **Mean** | **SD** |
| 5Hz | 13.20 | 0.51 |
| 1Hz | 13.08 | 0.44 |
| OFF | 13.02 | 0.40 |

Table S 13 Linear mixed model analysis results

LMM for duration, amplitude, frequency:

*[$metric] ~ Condition + (Condition|Subject) + (1|Electrode)*

**Duration (s)**

Summary:

| Term | F | df1 | df2 | p |
| --- | --- | --- | --- | --- |
| Intercept | 1347.10 | 1 | 1327 | <.001 |
| Condition | 1.49 | 2 | 1327 | .225 |

Fixed effects

| Factor | Estimate | SE | df | t | p | CI low | CI high |
| --- | --- | --- | --- | --- | --- | --- | --- |
| Intercept | 0.935 | 0.026 | 1327 | 36.70 | <.001 | 0.885 | 0.985 |
| 1 Hz | -0.015 | 0.032 | 1327 | -0.46 | .664 | -0.077 | 0.047 |
| 5 Hz | -0.057 | 0.034 | 1327 | -1.65 | .099 | -0.124 | 0.011 |

Random effects

| Factor | Variance | SD |
| --- | --- | --- |
| Subject (Intercept) | 0.096 | 0.310 |
| Subject (1 Hz) | 0.118 | 0.343 |
| Subject (5 Hz) | 0.134 | 0.366 |
| Electrode (Intercept) | 0.024 | 0.154 |
| Residual | 0.043 | 0.206 |

**Amplitude (μV)**

Summary

| Term | F | df1 | df2 | p |
| --- | --- | --- | --- | --- |
| Intercept | 310.27 | 1 | 1327 | <.001 |
| Condition | 0.59 | 2 | 1327 | 0.556 |

Fixed effects

| Factor | Estimate | SE | df | t | p | CI low | CI high |
| --- | --- | --- | --- | --- | --- | --- | --- |
| Intercept | 50.81 | 2.88 | 1327 | 17.61 | <.001 | 45.15 | 56.46 |
| 1 Hz | 1.40 | 1.70 | 1327 | 0.83 | 0.409 | -1.92 | 4.73 |
| 5 Hz | -0.20 | 1.09 | 1327 | -0.18 | 0.858 | -2.34 | 1.95 |

Random effects

| Factor | Variance | SD |
| --- | --- | --- |
| Subject (Intercept) | 8.578 | 2.929 |
| Subject (1 Hz) | 6.512 | 2.552 |
| Subject (5 Hz) | 3.541 | 1.882 |
| Electrode (Intercept) | 11.212 | 3.348 |
| Residual | 95.568 | 9.776 |

**Frequency (Hz)**

Summary

| Term | F | df1 | df2 | p |
| --- | --- | --- | --- | --- |
| Intercept | 22527.01 | 1 | 1327 | <.001 |
| Condition | 2.43 | 2 | 1327 | .089 |

Fixed effects

| Name | Estimate | SE | df | t | p | CI low | CI high |
| --- | --- | --- | --- | --- | --- | --- | --- |
| Intercept | 13.045 | 0.087 | 1327 | 150.09 | <.001 | 12.874 | 13.215 |
| 1 Hz | 0.032 | 0.093 | 1327 | 0.34 | .735 | -0.151 | 0.214 |
| 5 Hz | 0.152 | 0.079 | 1327 | 1.93 | .054 | -0.003 | 0.307 |

Random effects

| Random Effect | Variance | SD |
| --- | --- | --- |
| Subject (Intercept) | 0.289 | 0.538 |
| Subject (1 Hz) | 0.376 | 0.613 |
| Subject (5 Hz) | 0.320 | 0.565 |
| Electrode (Intercept) | 0.280 | 0.529 |
| Residual | 0.169 | 0.411 |

#### Spindle density (events/min)

Table S 14: Spindle density descriptives

| Condition | Mean (#/min) | SD |
| --- | --- | --- |
| 5Hz | 2.05 | 1.19 |
| 1Hz | 1.51 | 0.89 |
| Off | 1.44 | 0.93 |

#### Linear Mixed model analysis

Model equation:

*spindle_density ~ Condition + (Condition|Subject) + (1|Electrode)*

Table S 15: LMM model for spindle density

**Spindle density (#/min)**

Summary

| Term | F | df1 | df2 | p |
| --- | --- | --- | --- | --- |
| Intercept | 42.37 | 1 | 1797 | <.001 |
| Condition | 3.12 | 2 | 1797 | .044 |

Fixed effects

| Name | Estimate | SE | df | t | p | CI low | CI high |
| --- | --- | --- | --- | --- | --- | --- | --- |
| Intercept | 1.435 | 0.220 | 1797 | 6.51 | <.001 | 1.002 | 1.867 |
| 1 Hz | 0.075 | 0.174 | 1797 | 0.43 | .668 | -0.267 | 0.417 |
| 5 Hz | 0.617 | 0.272 | 1797 | 2.27 | .023 | 0.084 | 1.149 |

Random effects

| Random Effect | Variance | SD |
| --- | --- | --- |
| Subject (Intercept) | 0.887 | 0.942 |
| Subject (1 Hz) | 0.716 | 0.846 |
| Subject (5 Hz) | 1.174 | 1.084 |
| Electrode (Intercept) | 0.454 | 0.674 |
| Residual | 1.374 | 1.172 |

Post-hoc contrasts with Bonferroni-holm correction:

| Contrast | Estimate | t | p (uncorrected) | p (Holm) | Cohen’s d |
| --- | --- | --- | --- | --- | --- |
| 5Hz > 1Hz | 0.542 | 2.41 | .016 | .0481 | 0.334 |
| 5Hz > Off | 0.627 | 2.27 | .023 | .0481 | 0.380 |
| 1Hz > Off | 0.074 | 0.42 | .69 | .69 | 0.046 |

#### Spindle event density topography

Table S 16 Cluster corrected univariate test results:

Null distribution: 5000 subject-level sign-flip permutations, cluster threshold 0.15, minimum size = 2 electrodes

| **Contrast** | **Cluster mass value** | **Cluster, electrodes included (total = 30)** | **Cluster Mean** | **Cluster SD** | **p-value** | **Cohen’s d** |
| --- | --- | --- | --- | --- | --- | --- |
| **5Hz > 1Hz:** |  |  |  |  |  |  |
| Null, 95^th^ percentile | 23.82 |  | 1.64 | 0.98 |  |  |
| **Cluster 1** | 30.63 | 16/30 | 2.43 | 1.45 | .022 | 0.57 |
| **5Hz > OFF**: |  |  |  |  |  |  |
| Null, 95th percentile (OFF) | 23.49 |  | 1.32 | 0.97 |  |  |
| **Cluster 1** | 33.12 | 15/30 | 2.24 | 1.36 | .018 | 0.70 |
| **1Hz = OFF** |  |  |  |  |  |  |
|  |  |  |  |  | ns |  |

#### Spindle probability

The dependent variable was binary spindle occurrence (0/1) per trial (subject x electrode x trial x time bin). Total observations: 400,050; spindle events: 5,812 (1.45%).

##### Descriptive Statistics – Spindle Probability by Condition

Observed spindle probability per condition, computed as the mean of subject-level proportions.

Table S 17 Spindle probability descriptives

| Condition | Mean | SD |
| --- | --- | --- |
| OFF | 0.0125 | 0.0084 |
| 1 Hz | 0.0122 | 0.0065 |
| 5 Hz | 0.0171 | 0.0104 |

##### GLMM spindle probability per condition

Model used a binomial family with logit link, fitted via maximum likelihood (Laplace approximation).

M0 tests the overall effect of stimulation condition on spindle probability.

Table S 18 Model statistics

**Spindle probability**

Model

| Model | Formula | AIC | BIC |
| --- | --- | --- | --- |
| M0 | *Spindle ~ Condition + (1\|Subject) + (1\|Electrode)* | 59,402.41 | 59,456.90 |

Type III Wald Chi-Square Test

Effect	Chi-sq	df	p

Condition	40.64	2	< .001

Fixed Effects

(Estimates on the log-odds scale, sum-to-zero contrast coding was used)

| Parameter | Estimate | SE | z | p |
| --- | --- | --- | --- | --- |
| (Intercept) | -4.47 | 0.17 | -26.43 | < .001 |
| OFF | -0.06 | 0.02 | -2.87 | .004 |
| 1 Hz | -0.06 | 0.02 | -3.08 | .002 |

Post-Hoc Pairwise Contrasts (FDR-corrected)

| Contrast | OR | SE | z | p_FDR |
| --- | --- | --- | --- | --- |
| OFF / 1 Hz | 1.00 | 0.03 | 0.05 | .961 |
| OFF / 5 Hz | 0.84 | 0.03 | -5.24 | **< .001** |
| 1 Hz / 5 Hz | 0.84 | 0.03 | -5.54 | **< .001** |

Estimated Marginal Means (back transformed to probability).

| Condition | P(spindle) | SE | 95% CI |
| --- | --- | --- | --- |
| OFF | .0107 | .0018 | [.0077, .0149] |
| 1 Hz | .0107 | .0018 | [.0077, .0149] |
| 5 Hz | .0127 | .0021 | [.0092, .0177] |

##### Post stimulus-time histogram of spindle events

Table S 19 Spindle probaility descriptives

| Condition | Centre (s) | Mean | SD |
| --- | --- | --- | --- |
| OFF | -0.25 | 0.014 | 0.016 |
| OFF | 0.25 | 0.012 | 0.010 |
| OFF | 0.75 | 0.011 | 0.010 |
| OFF | 1.25 | 0.011 | 0.015 |
| OFF | 1.75 | 0.013 | 0.010 |
| 1 Hz | -0.25 | 0.011 | 0.009 |
| 1 Hz | 0.25 | 0.010 | 0.007 |
| 1 Hz | 0.75 | 0.011 | 0.009 |
| 1 Hz | 1.25 | 0.014 | 0.009 |
| 1 Hz | 1.75 | 0.015 | 0.014 |
| 5 Hz | -0.25 | 0.017 | 0.015 |
| 5 Hz | 0.25 | 0.017 | 0.013 |
| 5 Hz | 0.75 | 0.016 | 0.011 |
| 5 Hz | 1.25 | 0.018 | 0.018 |
| 5 Hz | 1.75 | 0.017 | 0.014 |

###### Post stimulus time histogram of spindle probability, GLMM model comparison

Five GLMMs of increasing complexity were compared.

All models used a binomial family with logit link, fitted via maximum likelihood (Laplace approximation). The best-fitting model (M5, bold) was selected based on AIC.

Table S 20: Model comparison results

| Model | Formula | AIC | BIC |
| --- | --- | --- | --- |
| M1 | *Spindle ~ Condition + TimeBin + (1\|Subject)* | 59,704.01 | 59,791.21 |
| M2 | *Spindle ~ Condition * TimeBin + (1\|Subject)* | 59,677.68 | 59,852.07 |
| M3 | *Spindle ~ Condition * TimeBin + (1\|Subject) + (1\|Electrode)* | 59,360.92 | 59,546.21 |
| M4 | *Spindle ~ Condition * TimeBin + (1\|Subject) + (1\|Subject:Electrode)* | 59,191.48 | 59,376.77 |
| **M5** | ***Spindle ~ Condition * TimeBin + (Condition\|Subject) + (1\|Electrode)*** | **59,090.33** | **59,330.12** |

Table S 21: Best Model (M5): Condition x Time Bin with Random Slopes

**Spindle probability**

Model summary - Type III Wald, Chi-Square Test

| Effect | Chi-sq | df | p |
| --- | --- | --- | --- |
| Condition | 8.09 | 2 | .018 |
| Time Bin | 22.06 | 4 | < .001 |
| Condition x Time Bin | 41.68 | 8 | < .001 |

Fixed Effects

Estimates on the log-odds scale. Sum-to-zero contrast coding was used.

| Factor | Estimate | SE | z | p |
| --- | --- | --- | --- | --- |
| (Intercept) | -4.53 | 0.18 | -25.85 | < .001 |
| Condition1 | -0.16 | 0.08 | -1.99 | .047 |
| Condition2 | -0.08 | 0.06 | -1.36 | .174 |
| TimeBin1 | -0.003 | 0.03 | -0.13 | .897 |
| TimeBin2 | -0.05 | 0.03 | -1.84 | .065 |
| TimeBin3 | -0.02 | 0.03 | -0.87 | .383 |
| TimeBin4 | -0.04 | 0.03 | -1.47 | .142 |
| Condition1:TimeBin1 | 0.13 | 0.04 | 3.32 | < .001 |
| Condition2:TimeBin1 | -0.05 | 0.04 | -1.31 | .191 |
| Condition1:TimeBin2 | 0.10 | 0.04 | 2.59 | .010 |
| Condition2:TimeBin2 | -0.17 | 0.04 | -4.33 | < .001 |
| Condition1:TimeBin3 | 0.01 | 0.04 | 0.33 | .739 |
| Condition2:TimeBin3 | 0.03 | 0.04 | 0.74 | .458 |
| Condition1:TimeBin4 | -0.14 | 0.04 | -3.48 | < .001 |
| Condition2:TimeBin4 | 0.11 | 0.04 | 2.84 | .004 |

Random Effects

| Factor | Parameter | SD |
| --- | --- | --- |
| Electrode | (Intercept) | 0.27 |
| Subject | (Intercept) | 0.74 |
| Subject | Condition1 | 0.32 |
| Subject | Condition2 | 0.22 |

Post-Hoc Contrasts – Conditions within Time Bins

Pairwise comparisons reported as odds ratios. p-values FDR-corrected within each time bin.

| Centre (s) | Contrast | OR | SE | z | p_FDR |
| --- | --- | --- | --- | --- | --- |
| -0.25 | OFF / 1 Hz | 1.11 | 0.14 | 0.78 | .433 |
| -0.25 | OFF / 5 Hz | 0.84 | 0.14 | -1.11 | .403 |
| -0.25 | 1 Hz / 5 Hz | 0.76 | 0.10 | -2.10 | .109 |
| 0.25 | OFF / 1 Hz | 1.22 | 0.16 | 1.52 | .130 |
| 0.25 | OFF / 5 Hz | 0.70 | 0.11 | -2.19 | **.043** |
| 0.25 | 1 Hz / 5 Hz | 0.58 | 0.08 | -4.11 | **< .001** |
| 0.75 | OFF / 1 Hz | 0.91 | 0.12 | -0.73 | .466 |
| 0.75 | OFF / 5 Hz | 0.72 | 0.12 | -2.03 | .110 |
| 0.75 | 1 Hz / 5 Hz | 0.79 | 0.11 | -1.79 | .110 |
| 1.25 | OFF / 1 Hz | 0.72 | 0.09 | -2.54 | **.016** |
| 1.25 | OFF / 5 Hz | 0.57 | 0.09 | -3.44 | **.002** |
| 1.25 | 1 Hz / 5 Hz | 0.79 | 0.10 | -1.80 | .072 |
| 1.75 | OFF / 1 Hz | 0.77 | 0.10 | -2.11 | .053 |
| 1.75 | OFF / 5 Hz | 0.61 | 0.10 | -3.07 | **.006** |
| 1.75 | 1 Hz / 5 Hz | 0.79 | 0.10 | -1.79 | .073 |

Note: OR < 1 indicates higher spindle probability for the second condition.

Effect Sizes – Odds Ratios for Fixed Effects

| Parameter | Log OR | SE | OR | 95% CI OR | z | p |
| --- | --- | --- | --- | --- | --- | --- |
| (Intercept) | -4.53 | 0.18 | 0.01 | [0.01, 0.02] | -25.85 | < .001 |
| Condition1 | -0.16 | 0.08 | 0.86 | [0.73, 1.00] | -1.99 | .047 |
| Condition2 | -0.08 | 0.06 | 0.93 | [0.83, 1.03] | -1.36 | .174 |
| TimeBin1 | -0.003 | 0.03 | 1.00 | [0.95, 1.05] | -0.13 | .897 |
| TimeBin2 | -0.05 | 0.03 | 0.95 | [0.90, 1.00] | -1.84 | .065 |
| TimeBin3 | -0.02 | 0.03 | 0.98 | [0.93, 1.03] | -0.87 | .383 |
| TimeBin4 | -0.04 | 0.03 | 0.96 | [0.91, 1.01] | -1.47 | .142 |
| Cond1:TimeBin1 | 0.13 | 0.04 | 1.14 | [1.05, 1.23] | 3.32 | **< .001** |
| Cond2:TimeBin1 | -0.05 | 0.04 | 0.95 | [0.88, 1.03] | -1.31 | .191 |
| Cond1:TimeBin2 | 0.10 | 0.04 | 1.11 | [1.03, 1.20] | 2.59 | .010 |
| Cond2:TimeBin2 | -0.17 | 0.04 | 0.84 | [0.78, 0.91] | -4.33 | **< .001** |
| Cond1:TimeBin3 | 0.01 | 0.04 | 1.01 | [0.94, 1.10] | 0.33 | .739 |
| Cond2:TimeBin3 | 0.03 | 0.04 | 1.03 | [0.95, 1.11] | 0.74 | .458 |
| Cond1:TimeBin4 | -0.14 | 0.04 | 0.87 | [0.80, 0.94] | -3.48 | **< .001** |
| Cond2:TimeBin4 | 0.11 | 0.04 | 1.11 | [1.03, 1.20] | 2.84 | .004 |

Estimated Marginal Means

(EMMs and CIs are back transformed to probability)

| Condition | Centre (s) | P(spindle) | SE | 95% CI |
| --- | --- | --- | --- | --- |
| OFF | -0.25 | .010 | .002 | [.007, .016] |
| OFF | 0.25 | .010 | .002 | [.006, .015] |
| OFF | 0.75 | .009 | .002 | [.006, .014] |
| OFF | 1.25 | .008 | .002 | [.005, .012] |
| OFF | 1.75 | .009 | .002 | [.006, .014] |
| 1 Hz | -0.25 | .009 | .002 | [.007, .013] |
| 1 Hz | 0.25 | .008 | .001 | [.006, .011] |
| 1 Hz | 0.75 | .010 | .002 | [.007, .014] |
| 1 Hz | 1.25 | .011 | .002 | [.007, .015] |
| 1 Hz | 1.75 | .012 | .002 | [.009, .017] |
| 5 Hz | -0.25 | .012 | .002 | [.009, .017] |
| 5 Hz | 0.25 | .014 | .002 | [.010, .019] |
| 5 Hz | 0.75 | .013 | .002 | [.009, .018] |
| 5 Hz | 1.25 | .013 | .002 | [.009, .019] |
| 5 Hz | 1.75 | .015 | .003 | [.011, .022] |
